# Brain connectivity-based prediction of real-life creativity is mediated by semantic memory structure

**DOI:** 10.1101/2021.07.28.453991

**Authors:** Marcela Ovando-Tellez, Yoed N. Kenett, Mathias Benedek, Matthieu Bernard, Joan Belo, Benoit Beranger, Theophile Bieth, Emmanuelle Volle

**Affiliations:** Sorbonne University, FrontLab at Paris Brain Institute (ICM), INSERM, CNRS, 75013, Paris, France; Faculty of Industrial Engineering and Management, Technion – Israel Institute of Technology, Haifa 3200003 Israel; Institute of Psychology, University of Graz, Graz, Austria; Sorbonne University, CENIR at Paris Brain Institute (ICM), INSERM, CNRS, 75013, Paris, France; Neurology department, Pitié-Salpêtrière hospital, AP-HP, F-75013, Paris, France

**Keywords:** creativity, semantic network, brain networks, functional connectivity, cognition

## Abstract

Creative cognition relies on the ability to form remote associations between concepts, which allows to generate novel ideas or solve new problems. Such an ability is related to the organization of semantic memory; yet whether real-life creative behavior relies on semantic memory organization and its neural substrates remains unclear. Therefore, this study explored associations between brain functional connectivity patterns, network properties of individual semantic memory, and real-life creativity. We acquired multi-echo functional MRI data while participants underwent a semantic relatedness judgment task. These ratings were used to estimate their individual semantic memory networks, whose properties significantly predicted their real-life creativity. Using a connectome-based predictive modeling approach, we identified patterns of task-based functional connectivity that predicted creativity-related semantic memory network properties. Furthermore, these properties mediated the relationship between functional connectivity and real-life creativity. These results provide new insights into how brain connectivity supports the associative mechanisms of creativity.

## Introduction

Creativity is key to our ability to cope with change, innovate, and find new solutions to address societal challenges (*1*). Understanding the complex and multidimensional construct of creativity is thus fundamental to support societal, cultural, and economic progress. Creative behaviors in real life depend on individual differences in cognitive ability, in addition to personality and environmental factors (*2*). The cognitive mechanisms underlying creative abilities are not yet understood (*3*–*6*). The associative theory hypothesizes that creative abilities are related to the organization of semantic associations in memory (*7*). In support of this theory, several studies found that more creative individuals are able to link distant concepts more easily (*8*–*10*), have less common or constrained word associations, and a more flexible organization of semantic memory (*9*, *11*–*15*). In addition, in brain-damaged patients, rigid semantic associations were associated with poor creative abilities (*16*–*18*). Associative thinking has been related to creative abilities as measured within several existing frameworks, such as divergent thinking (*8*, *9*, *14*, *19*–*21*), insight problem solving (*7*, *22*), analogical reasoning (*23*, *24*), as well as to creative achievements in real life (*25*–*28*). Overall, the properties of semantic memory play an essential role in the cognitive processes that bring forth original ideas.

Recent research has demonstrated how computational network science methodologies (*29*–*32*) based on mathematical graph theory allow exploring the properties and organization of the concepts in semantic memory via semantic networks (SemNets). Applying these methods, several studies have shown that creative abilities can be related to semantic memory organization (*11*, *33*–*38*). Kenett and colleagues (*11*) investigated the SemNets of groups of low and high creative individuals, based on free associations generated by both groups to a list of 96 cue words. They found that the SemNets of low creative individuals were less connected and more spread out compared to the SemNets of high creative individuals. However, estimating SemNets at the group level may obscure individual differences related to creativity. To address this issue, Benedek and colleagues (*36*) developed a method to estimate individual SemNets, based on word relatedness judgment ratings. Participants rated the relationships between all possible pairs of 28 cue words, serving as a proxy for the organization of these words in an individuals’ semantic memory. They demonstrated how individual-based SemNet metrics replicated the group-based findings of Kenett et al. (*11*), and were related to individual differences in divergent thinking scores (the most widely assessed component of creative thinking) (*39*, *40*). A recent study reported similar results (*41*). In a previous study, (*37*) we replicated and extended this finding with two improvements: We controlled the selection of the cue words using a computational method optimizing the distribution of theoretical distances between words, and we assessed creative abilities and behaviors using a more diverse set of tools. This study showed that the network metrics of the individual SemNets correlated with several measures of creativity, including a questionnaire of creative activities and achievements (*42*). Hence, individual SemNets measures—reflecting the properties of semantic memory—allow exploring underlying cognitive mechanisms of creativity, suggesting that more creative individuals have more flexible semantic associations and connect more distant concepts or words (*38*). However, the neurocognitive determinants of individual differences in creativity related to the flexibility of semantic associations are still unclear and unexplored.

Existing MRI-based neuroimaging studies have identified a large set of brain regions involved in creative cognition (*5*, *12*, *43*–*47*). A growing body of creativity neuroscience research has highlighted the importance of functional interactions within and between several brain networks, including the executive control network, salience network and the default mode network (*5*, *48*). Additionally, semantic and episodic memory regions (*44*, *49*–*52*) and the motor and premotor regions have been shown to play a role in creative cognition (*44*, *53*). The advantage of a whole-brain functional connectivity approach is to provide a holistic and functional view of how brain networks relate to creative thinking. For example, resting-state functional connectivity within and between these networks was shown to predict creative abilities (*54*, *55*) and task-based functional connectivity within and between these networks increased during a creativity task, compared to a control task (*5*, *43*). A recent approach in neuroimaging research is connectome-based predictive modeling (CPM) (*56*), which uses machine learning methods to identify patterns of functional connectivity that predict complex cognitive functions, including divergent thinking ability (*43*, *56*–*61*). Unlike previous research that focused on the brain connectivity associated with specific creativity tasks (e.g., divergent thinking), the current study explores the neurocognitive determinants of real-life creativity by studying the neural basis of semantic memory organization related to creative behavior. We hypothesized that the associative mechanisms reflected by SemNet metrics are relevant to real-life creative activities and achievements and can be predicted by functional connectivity patterns, involving, in particular, the control, default, and salience networks (*43*).

To this end, we first examine the organization of individual SemNets via network metrics and identify the SemNet metrics that reliably predict differences in creative achievement and thus constitute cognitive markers of real-life creativity. We then explore the functional connectivity of brain networks predicting individual differences in these SemNet markers. We use the CPM method and analyze functional brain connectivity during the performance of the semantic relatedness task that is used to estimate individual SemNets. We identify the task-based functional connectivity patterns predicting individual differences in SemNet properties. Finally, we examine whether SemNet properties mediate the link between these brain connectivity patterns and real-life creativity, thus linking functional connectivity to real-life creativity via individual differences in semantic memory organization.

## Results

### Individual Semantic Network metrics and creativity

First, we explored the properties of individuals’ SemNets in relation to creativity. Similar to previous studies (*36*–*38*), we estimated participants’ individual semantic memory network as weighted (WUN) and unweighted (UUN) SemNets based on performance in the semantic relatedness judgment task (RJT; **Figure 1**). During the RJT, participants judged the relatedness between all possible pairs of 35 words (595 ratings). We then computed established network measures in cognitive network research including (*29*): Average Shorter Path Length (*ASPL;* measuring average distances, or the spread of the SemNet), Clustering Coefficient (*CC;* measuring overall connectivity in the SemNet), Modularity (*Q;* measuring the level of segregation of the SemNet) and Small Worldness (*S;* measuring the ratio between connectivity and distances in the network (*62*), see *Material and Methods*). In addition, we assessed individual differences in real-life creative activities (*C-Act*) and achievements (*C-Ach*) via the Inventory of Creative Activities and Achievements (*42*) completed outside the MRI scanner (Descriptive statistics for behavioral and network measures are reported in **Table 1**).

**Table 1.** Descriptive statistics of creativity scores and semantic network measures. Data are shown for real-life creativity activities (*C-Act*) and achievements (*C-Ach*), and for SemNet metrics of weighted (WUN) and unweighted (UUN) networks.

|  | *Mean* | *SD* | *Min* | *Max* |
| --- | --- | --- | --- | --- |
| **Creativity scores** |  |  |  |  |
| *C-Act* | 47.894 | 21.695 | 13 | 102 |
| *C-Ach* | 74.638 | 42.249 | 1 | 207 |
| **WUN metrics** |  |  |  |  |
| *ASPL* | 0.021 | 0.004 | 0.015 | 0.037 |
| *CC* | 0.363 | 0.096 | 0.142 | 0.628 |
| *Q* | 0.122 | 0.058 | 0.032 | 0.319 |
| *S* | 1.003 | 0.073 | 0.828 | 1.387 |
| **UUN metrics** |  |  |  |  |
| *ASPL* | 1.633 | 0.221 | 1.262 | 2.361 |
| *CC* | 0.585 | 0.082 | 0.438 | 0.781 |
| *Q* | 0.178 | 0.064 | 0.058 | 0.392 |
| *S* | 1.386 | 0.271 | 1.011 | 2.936 |

Note. *ASPL*= Average Shortest Path Length; *CC* = Clustering coefficient; *Q* = Modularity; *S* = Small-Worldness.

**Figure 1.**
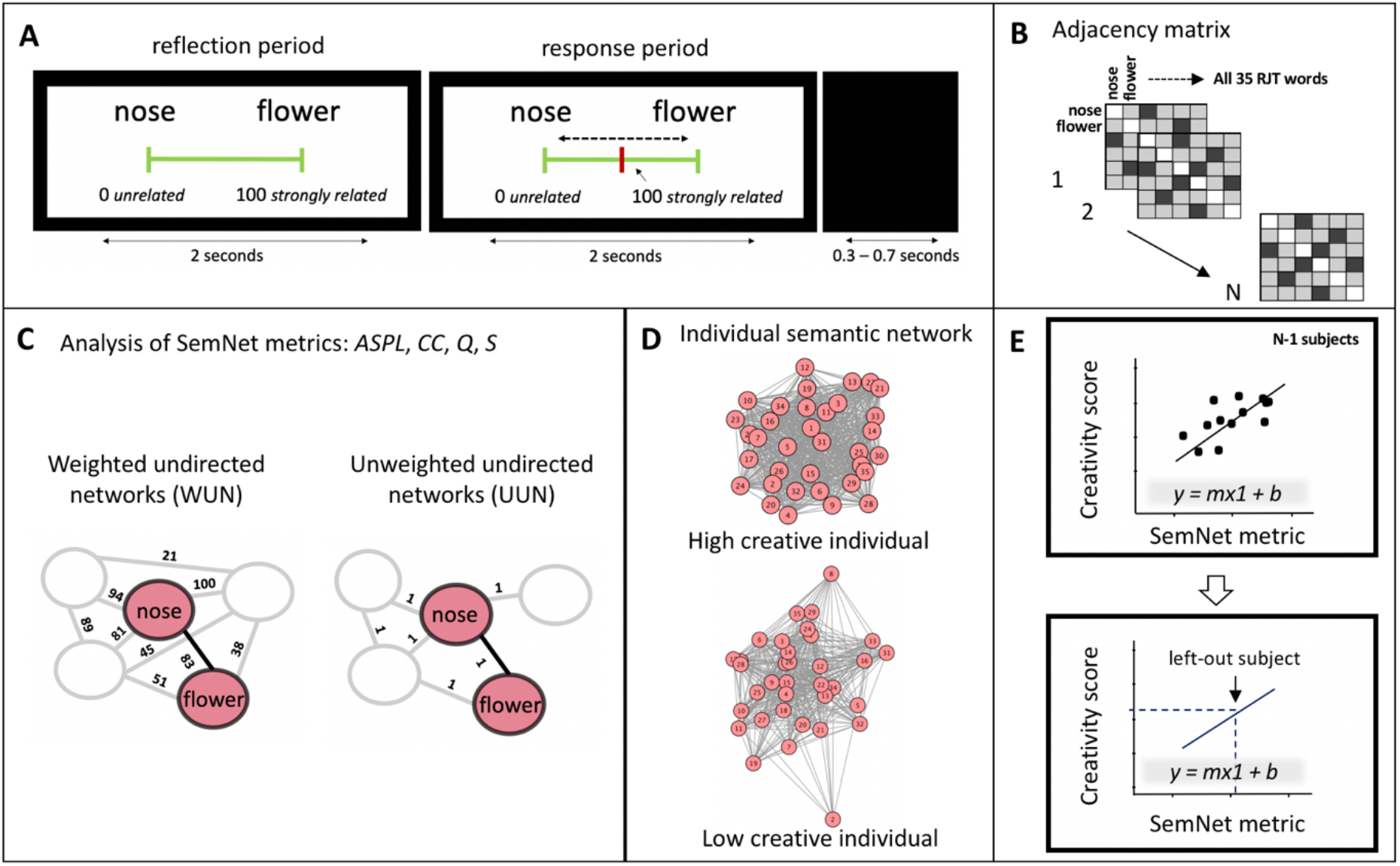
Estimation of individual semantic networks (SemNets) to predict creativity. (**A**) Trial representation of an exemplary trial of the RJT asking participants to judge the relatedness of 595 word pairs. Each trial began with the display of a pair of words along with a visual scale (reflection period) ranging from 0 (unrelated words) to 100 (strongly related words). During the next 2 seconds (response period), participants were allowed to move the cursor (in red) using a trackball to indicate the relatedness of the two words. An intertrial interval of 0.3-0.7s separated trials. (**B**) For each participant, we computed a 35 by 35 adjacency (connectivity) matrix with columns and rows representing each of the 35 RJT words, and cell values correspond to the relatedness judgments given by the participant during the RJT. (**C**) We estimated individual semantic memory networks following two established approaches: weighted (WUN) and unweighted (UUN) undirected networks, using the RJT words as the network nodes. In the WUN networks, the RJT judgments reflected the strength of links between nodes. In the UUN networks, the RJT judgments above average (50) were kept and set to one. The SemNet metrics were computed for both WUN and UUN separately: *ASPL*, *CC*, *Q* and *S*. (**D**) Representation of the individual WUN SemNets for a low creative and a high creative participant. (**E**) Linear regressions using leave-one-out cross-validations were performed to explore whether real-life creative activities (*C-Act*) and achievements (*C-Ach*) were predicted from SemNet properties estimated in (b). The SemNet metrics were used to build predictive linear models in N-1 participants. The predictive model was tested on the left-out participant using its SemNet metric (m) to predict its creativity scores. RJT = relatedness judgment task; SemNet = semantic network; ASPL = average shortest path length; CC = clustering coefficient; Q = modularity; S = small-worldness.

We then examined how SemNet metrics predict real-life creativity by applying linear regression models, regressing creativity on each SemNet metric with leave-one-out cross-validations: We iteratively fitted predictive linear models in N-1 participants and tested the model in the left-out participant. The significance of the model prediction was assessed by the correlation between the predicted value of *C-Act* (or *C-Ach*) computed by the model and the observed value using permutation testing. These analyses revealed that both real-life creative activities and achievements are predicted from different individual SemNet metrics (**Figure 2**). The Spearman correlations showing the direction and size of the relationships between SemNet metrics and creativity are reported in **Table 2**. *C-Act* was predicted from WUN *ASPL* and UUN *Q. C-Ach* was predicted from WUN *Q* and UUN *Q*. More creative individuals had less modular SemNets.

**Table 2.** Relationship between individual semantic network metrics and creativity. The Spearman correlations between SemNet metrics and creativity scores are reported (*r_s_* for *C-Act* and *C-Ach*). In bold are the significant predictions of creativity from the SemNet properties after permutation testing shown in **Figure 2**. * indicate correlations that reached significance after FDR correction for multiple comparisons.

| Creativity scores | *C-Act* |  | *C-Ach* |  |
| --- | --- | --- | --- | --- |
| | $r_s$ | $p$ | $r_s$ | $p$ |
| **WUN metrics** |  |  |  |  |
| *ASPL* | **-.276** | **.007\*** | -.208 | .044 |
| *CC* | .165 | .111 | .201 | .052 |
| *Q* | -.179 | .085 | **-.295** | **.004\*** |
| *S* | .234 | .023 | -.017 | .868 |
| **UUN metrics** |  |  |  |  |
| *ASPL* | -.125 | .230 | -.149 | .152 |
| *CC* | .092 | .378 | .080 | .441 |
| *Q* | **-.281** | **.006\*** | **-.287** | **.005\*** |
| *S* | -.154 | .139 | -.219 | .034 |

**Figure 2.**
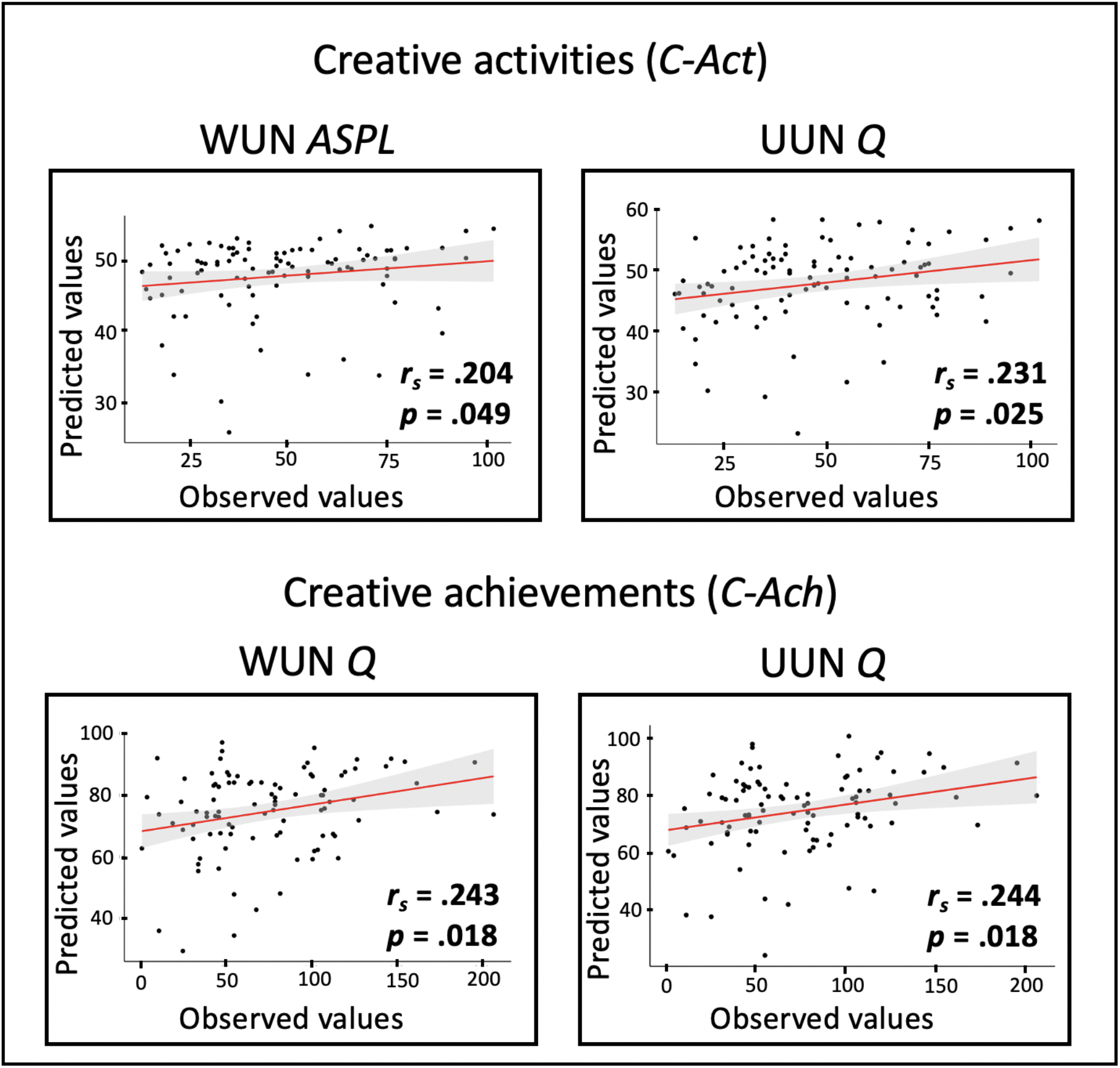
Prediction of creativity scores from Semantic network metrics. The plots show the Spearman correlations between the predicted values (y-axis) and observed values (x-axis) of creative activities and achievements based on individual SemNet metrics for the significant predictions. At the bottom-right part of each plot, we present the *r*_s_ and the *p* values, based on permutation testing.

### Prediction of creativity-related SemNet properties from brain connectivity

We applied the connectome-based predictive modeling (CPM) approach (*43*, *56*, *57*, *59*) to explore whether task-based functional connectivity patterns predict semantic memory network metrics that related to creativity (i.e., *Q* in WUN and UUN, and *ASPL* in WUN; see **Table 2**; The applied CPM approach is illustrated in **Figure 3**). We used a functional brain atlas to define 200 brain nodes belonging to 17 functional networks (*63*). For each participant, Pearson correlations of the BOLD signal between all unique pairs of brain regions (i.e., nodes; n = 19,900) were computed to estimate the task-related functional connectivity of the whole brain connectivity network (**Figure 3a**). We then identified relevant links of the brain connectivity network that positively (positive model network) or negatively (negative model network) correlated with the SemNet metric across participants (**Figure 3b**). Next, we adapted the classical CPM method (*56*) to better take into account the network properties of the brain model networks. Instead of using the sum of the connectivity in the model networks, we computed two key network metrics describing small-worldness properties of human brain networks (*64*–*66*): their *CC* (*brain-CC*) and *efficiency* (*brain-Eff*; **Figure 3c**). We then ran six separate linear models regressing each SemNet metric (*Q* for WUN and UUN, and *ASPL* for WUN) on each model network metric (*brain-CC* and *brain-Eff*). We used leave-one-out cross-validations, iteratively fitting predictive linear models in N-1 participants and tested these models on the left-out participant (**Figure 3d**). Finally, the model prediction was assessed by the Spearman correlation between the predicted value from the model and the observed values.

**Figure 3.**
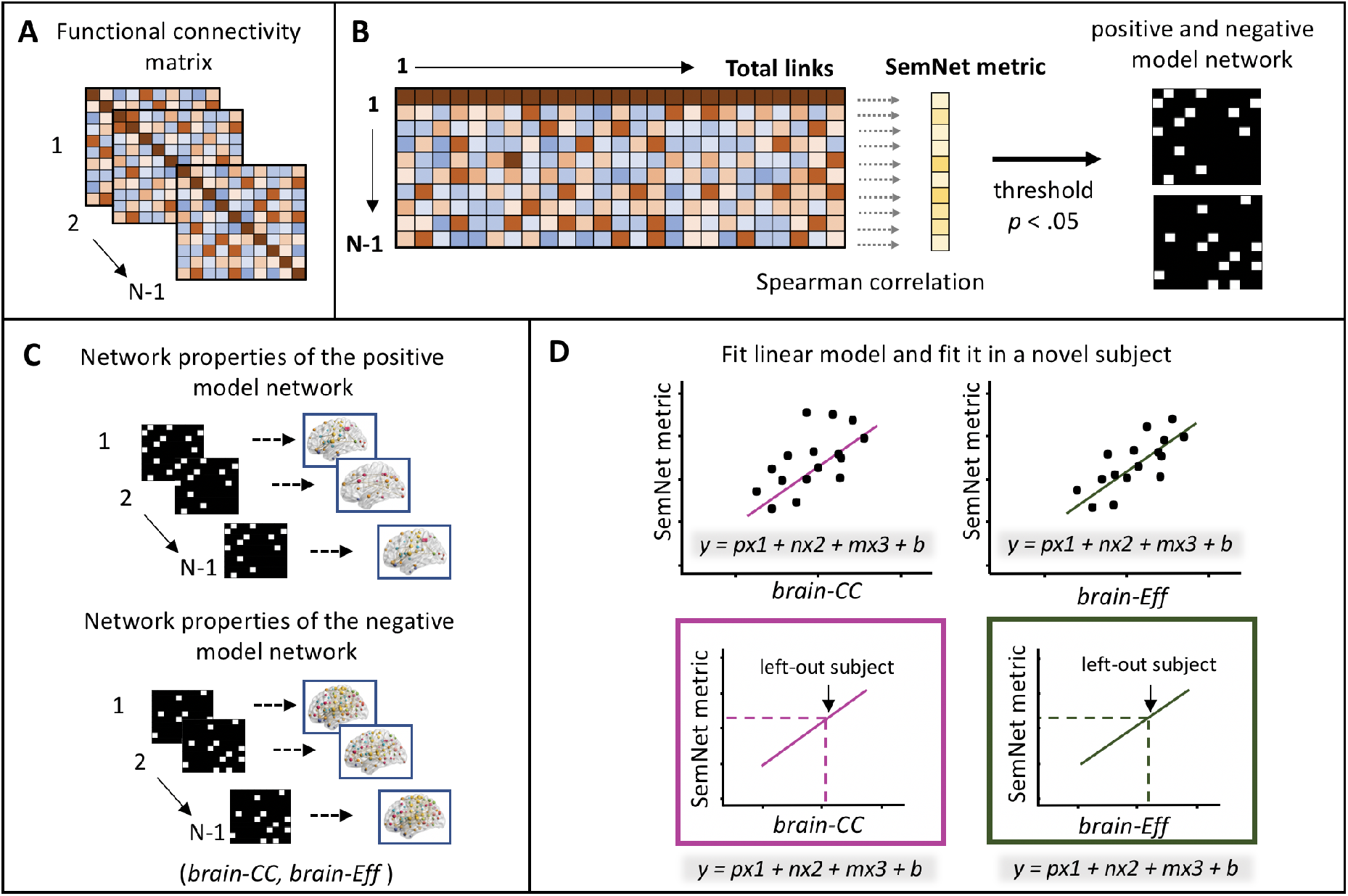
Connectome Predictive Modeling-based prediction method. (**A**) We defined the brain nodes based on the Schaefer atlas consisting of 200 ROIs (*63*). For each participant, we assessed the BOLD activity during the RJT in each ROI and used pairwise Pearson correlations to estimate a 200 by 200 task-related functional connectivity matrix. Using a leave-one-out approach, all of the CPM steps were conducted in N-1 participants. (**B**) The functional connectivity matrix (all links) was correlated to SemNet metrics using Spearman correlations. The links that significantly positively or negatively correlated with the SemNet metric (*p* < .05) formed a positive and a negative model network, respectively. (**C**) We calculated two network properties (in separate CPM analyses) of the positive and negative model networks, *brain-CC* and *brain-Eff* metrics. (**D**) The brain metrics in the positive (p) and negative (n) model networks were used to build a linear model predicting the SemNet metric in the left-out participant. Since head motion can impact CPM, we included the meanFD variable (m), a head motion parameter, as a regressor in the model to avoid a possible effect in the prediction. Finally, the model was applied to the left-out participant to compute a predicted SemNet value from his/her brain model networks. The predicted value was then correlated with the observed value to assess the model predictive validity.

We then tested the relation between predicted and observed CPM models on the various SemNet metrics, using 1,000 iteration permutation testing (*56*) (**Figure 4**). The CPM-based prediction from *brain-CC* was significant for the WUN *Q* metric (*r* = .386, *p* = .004). The CPM-based predictions from *brain-Eff* were significant for the WUN *Q* metric (*r* = .476, *p* = .001) and the UUN *Q* metric (*r* = .272, *p* = .036). The CPM-based predictions of WUN *ASPL* from both *brain-CC* and *brain-Eff*, and UUN *Q* from *brain-CC* were not significant, showing either a negative correlation between predicted and observed values or did not reach a significant *p*-value after permutation testing. In summary, CPM analyses on task-based functional connectivity showed that brain connectivity *CC* and *efficiency* allowed reliable predictions of SemNet *Q*.

**Figure 4.**
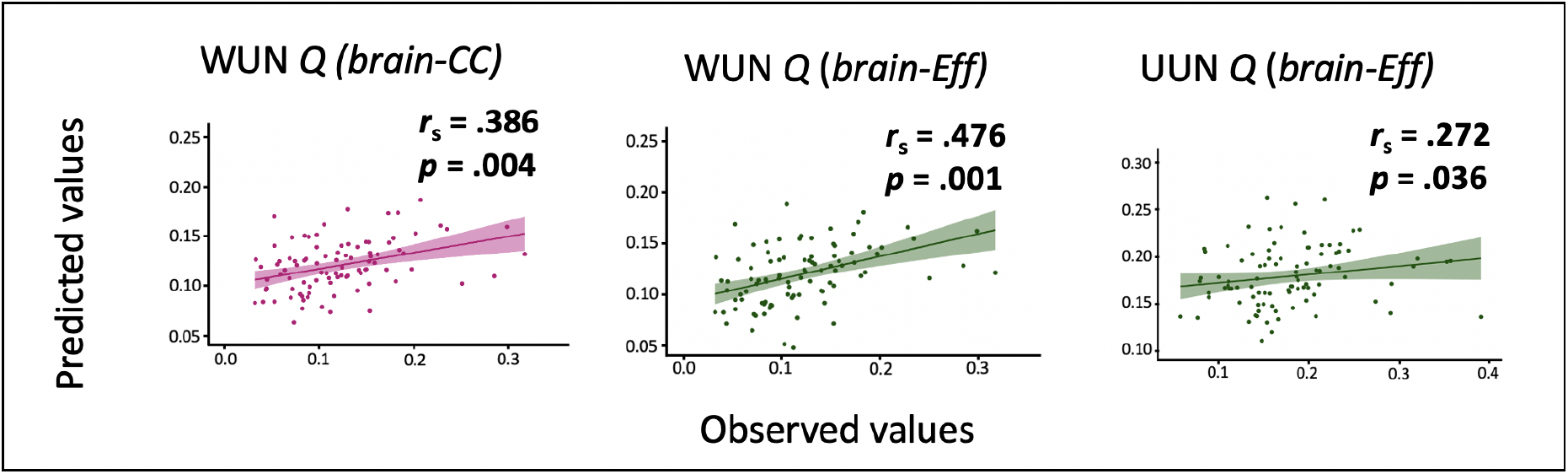
Predicted and observed SemNet metrics. The plots show the Spearman correlations between the predicted values (y-axis) and observed values (x-axis) of SemNet metrics based on brain connectivity for the significant predictions. Green plots are presented for *brain-Eff* and magenta ones for *brain-CC.* In the upper-right side of each plot, we present the *r*_s_ and the *p* values. The reported *p* values are based on permutation testing.

### Functional anatomy of the predictive brain connectivity patterns

To characterize the functional brain connectivity patterns predictive of SemNet metrics, we explored the links of the model networks that account for SemNet properties relevant to creativity. Unique positive and negative model networks were identified for each SemNet metric (*56*) (**Figure 3b**) and used to compute their network properties (*brain-CC* and *brain-Eff*; **Figure 3c**). Since SemNet *modularity* (*Q*) was negatively correlated with both creativity measures (*C-Act* and *C-Ach*; **Table 2**) as expected from previous studies (*11*, *36*–*38*), we focused on the description of the negative model network predicting *UUN Q* **(Figure 5)** or WUN *Q* (**SI Figure S1**). In this model network, we considered the links that were shared in all iterations of the leave-one-out analysis, as the links in the model network can slightly vary at each iteration.

**Figure 5.**
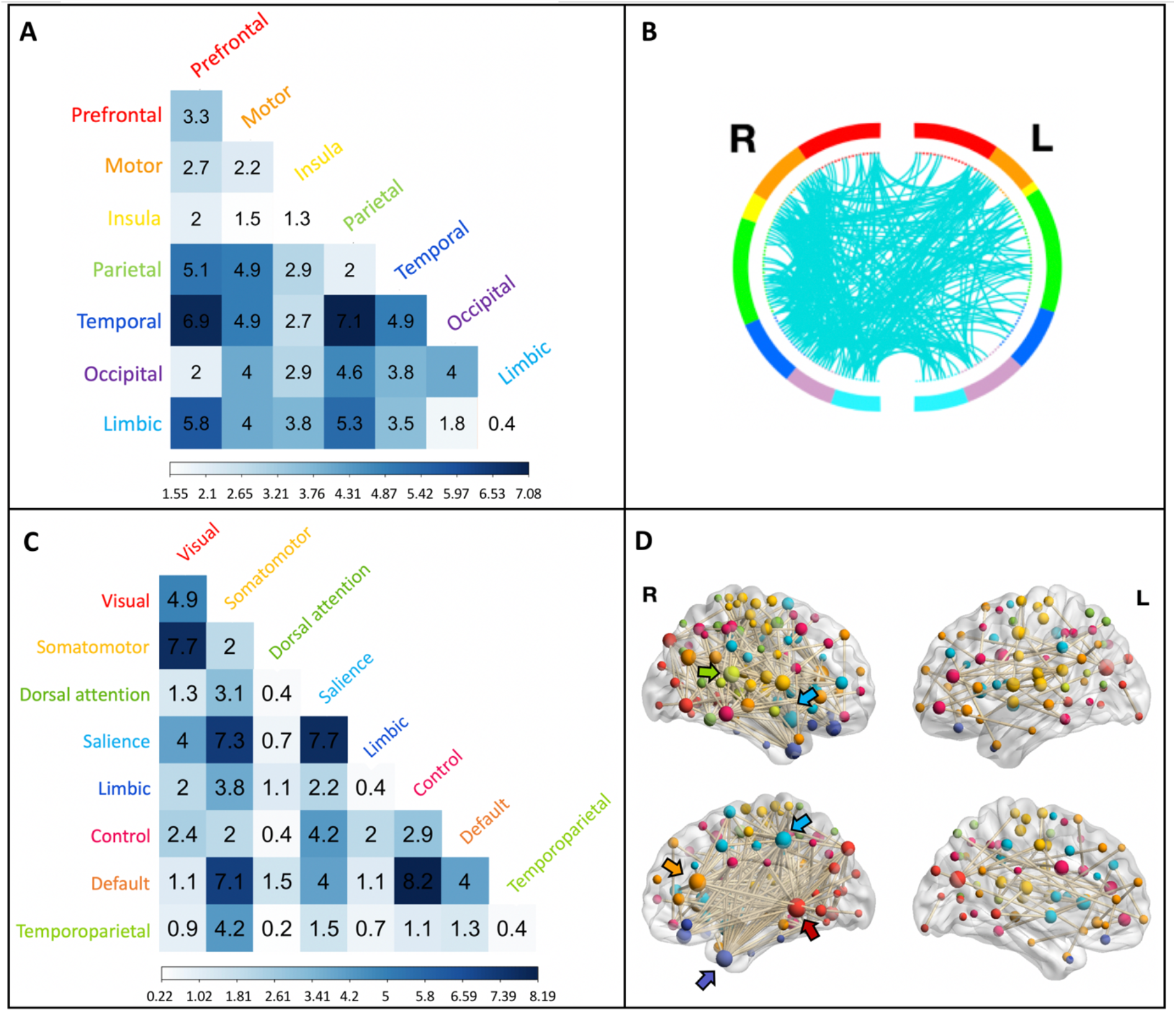
Functional anatomy of the CPM model predicting the SemNet metric UUN *Q*. (**A**) First, we examined the distribution of the links of the model network at the brain location level, specifically into the brain lobes. The correlation matrix represents the percentage of links within the model network connecting seven different brain lobes (total links = 452). (**B**) A circular graph represents the distribution of links within and between brain regions in the left and right hemispheres. Brain regions are color-coded as in (**A**), and the cyan lines represent the links connecting the ROIs. For visualization purposes, we used a nodal degree threshold of *k* > 10. (**C**) Second, we examined the distribution of the links across intrinsic functional networks based on Schaefer’s atlas *(63)*. The matrix represents the percentage of links within the model network occurring within and between eight intrinsic brain networks. (**D**) The nodes and links of the model network are superimposed on a volume rendering of the brain. The color of the nodes represents the functional network they belong to, using a similar color code as in (**B**). The size of the nodes is proportional to their degree, and the highest degree nodes are marked by arrows. Nodes with degree *k* = 0 are not displayed.

For the standard CPM negative model network of UUN *Q*, we identified 452 links. Connectivity of these links related to lower SemNet *Q*, which again predicted higher real-life creativity. These links represented connections mainly within and between temporal, parietal, limbic and prefrontal lobes (**Figure 5a–b**). When we explored the distribution of these links at the functional networks level, based on the functional networks included in the Schaefer atlas (*63*), most of the links were part of the somatomotor, salience and default mode networks (**Figure 5c**). The highest number of links were found between control and default mode networks (8.2%), followed by links within the salience network and between somatomotor and visual networks. In this model network, the highest degree nodes — nodes with highest number of connections (*k*; i.e., the number of functional connections) — belonged to the right hemisphere being part of the visual network (i.e., extra-striate inferior, *k* = 53), default mode network (i.e., medial prefrontal cortex, *k* =39), salience (i.e., insula, *k* = 31; parietal medial, *k* = 28), temporoparietal (i.e., temporal-parietal; *k* =29) and limbic (temporal pole, *k* = 28) networks (**Figure 5d**). In summary, the main patterns of functional connectivity that predicted lower SemNet *Q* (i.e., related to higher creativity) had a whole-brain distribution and involved the control, default mode, salience and somatomotor networks.

### Mediation Analysis

In the previous analyses, we found a relationship between SemNets and real-life creativity, and between brain functional connectivity and SemNets. In a final step, we analyzed whether the relationship between functional brain connectivity and real-life creativity is mediated by the SemNet properties. Hence, we conducted mediation analyses that focused on the indirect effect of functional connectivity on creative activities and achievements, using either *C-Act* or *C-Ach* as the dependent variable for each significant CPM model. To simplify interpretations, since UUN *Q* had a negative correlation with creativity, its value was reversed (UUN *Q*R) to be positively correlated with creativity.

Since *C-Act* was significantly predicted by the SemNet metric UUN *Q,* we explored the mediating role of UUN *Q* on the relationship between the properties of the functional brain network predicting UUN *Q* (*brain-Eff*) and *C-Act* (**Figure 6a**). As shown in the previous analyses, the regression coefficient between *brain-Eff* and *UUN Q*R was statistically significant (*beta* = .305, *p* < .001), as was the regression coefficient between *UUN Q*R and *C-Act* (*beta* = .443, *p* = .002). The total effect and the direct effect were not statistically significant (*beta* = .116, *p* = .328; *beta* = −.019, *p* = .872). We tested the significance of the indirect effect using a bootstrapping method. The bootstrapped indirect effect was (.305)*(.443) = .135, and the 95% confidence interval ranged from 0.024 to 0.320. Thus, the indirect effect was statistically significant (*p* = .002). Hence, SemNets UUN *Q* mediated the relationship between the efficiency of functional brain connectivity (*brain-Eff*) and creative activities (*C-Act*): The higher the efficiency of the negative model network that predicts UUN *Q*, the lower the SemNet *Q*, and the higher are real-life creative activities.

**Figure 6.**
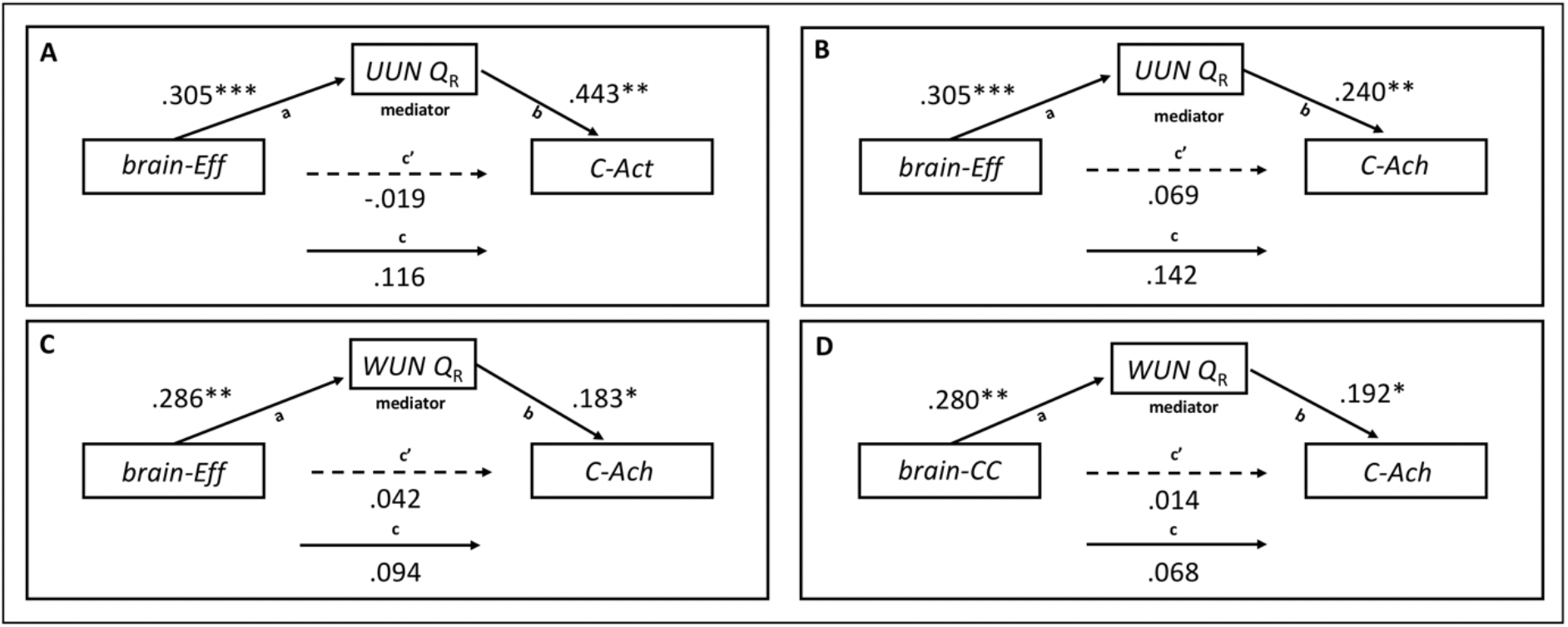
Mediation Analyses. Results of the mediation models are presented in path diagrams. Each diagram indicates the beta weights of the regression coefficients with the brain metrics of the model network (*brain-Eff* and *brain-CC*) as the independent variable (predictor), SemNet metrics as the mediator (*UUN Q*R and *WUN Q*R), and real-life creativity (*C-Act* and *C-Ach*) as the dependent variable (outcome). The total effect is indicated by path c, the direct effect by path c’, and the indirect effect is given by the product of path a and path b. The indirect effect was significant in all the reported mediations (**A**) The mediating role of UUN *Q* on the relationship between the *brain-Eff* of the brain functional network predicting it and *C-Act.* (**B**) Mediating role of UUN *Q* between the *brain-Eff* of the brain network predicting it and *C-Ach.* (**C**) Mediating role of the weighted networks WUN *Q* on the relationship between the *brain-Eff* of the functional connectivity of the negative model network predicting it and *C-Ach.* (**D**) Mediating role of WUN *Q* on the relationship between the *brain-CC* of the functional connectivity of the negative model network predicting it and *C-Ach.* * *p* < .05; ** *p* < .01; *** *p* < .001

*C-Ach* score was predicted from SemNet WUN *Q* and UUN *Q* metrics. We explored the mediating role of UUN *Q* between the functional connectivity of the negative model network predicting it (*brain-Eff*) and *C-Ach* (**Figure 6b**). The mediation analysis showed that the regression coefficient between *brain-Eff* and *UUN Q* R was statistically significant (*beta* = .305, *p* < .001), as was the regression coefficient between the *C-Ach* and *UUN Q*R (*beta* = .241, *p* = .005). The total effect and the direct effect were not statistically significant (*beta* = .142, *p* = .056; *beta* = .069, *p* = .353). The bootstrapped indirect effect was (.305)*(.241) = .073, and the 95% confidence interval ranged from 0.018 to 0.140. Thus, the indirect effect was statistically significant (*p* < .001).

Hence, SemNets UUN *Q* mediated the link between the efficiency of brain functional connectivity (*brain-Eff*) and real-life creative achievements (*C-Ach*): The higher the efficiency of the negative model network that predicts UUN *Q*, the lower the *modularity* of SemNet, and the higher the real-life creative achievements.

Similarly, we explored the mediating role of WUN *Q* on the relationship between the properties of the functional connectivity of the negative model network predicting it *(brain-Eff and brain-CC)* and *C-Ach* (**Figure 6c**). Using *brain-Eff* as an independent variable, the regression coefficient between *brain-Eff* and *WUN Q*R was significant (*beta* = .286, *p* = .004), as was the regression coefficient between *C-Ach* and *WUN Q*R (*beta* = .183, *p* = .015). The total effect and the direct effect were not statistically significant (*beta* = .094, *p* = .183; *beta* = .042, *p* = .560). The bootstrapped indirect effect was (.286)*(.183) = .052, and the 95% confidence interval ranged from 0.005 to 0.110. Thus, the indirect effect was statistically significant (*p* = .018).

Using *brain-CC* as independent variable, the regression coefficient between *brain-CC* and *WUN Q*R was significant (*beta* = .280, *p* = .008), as was the regression coefficient between *C-Ach* and *WUN Q*R (*beta* = 0.192, *p* = .01) (**Figure 6d**). The total effect and the direct effect were not statistically significant (*beta* = .068, *p* = .365; *beta* = .014, *p* = .850). The bootstrapped indirect effect was (.280)*(.192) = .054, and the 95% confidence interval ranged from 0.006 to 0.130. Thus, the indirect effect was statistically significant (*p* = .018).

Hence, SemNet WUN *Q* mediated the link between the efficiency (*brain-Eff*) and the clustering coefficient (*brain-CC*) of functional brain connectivity and real-life creative achievements (*C-Ach*): The higher the efficiency and clustering of the negative model network that predicted WUN *Q*, the lower SemNets *Q*, and the higher the real-life creative achievements. In summary, individual SemNets *Q* measured in WUN and UUN networks mediated the relationship between brain functional connectivity and real-life creativity.

## Discussion

Our results provide a new neuroscientific understanding of the individual determinants of real-life creative behavior. Recently developed computational approaches allowed us to predict complex cognitive functions from brain connectivity (*56*–*58*) and to explore the organization of semantic memory at the individual level using SemNets (*36*–*38*). The unprecedented combination of these approaches revealed unique patterns of brain functional connectivity that reliably predict differences in real-life creativity via semantic network structure. Using the CPM approach, we show that brain connectivity during semantic relatedness judgments predicted individual differences in the *modularity* (*Q*) of SemNets that was identified as a behavioral marker of creativity. Specifically, the efficiency and clustering of whole-brain connectivity patterns predicted differences in real-life creativity mediated by SemNet *modularity*.

According to the associative theory of creativity ^7^, high creative individuals are characterized by a more flexible organization of concepts in their semantic memory, allowing them to retrieve remote associations more easily (*14*, *16*). A recent study revealed the mediating role of associative abilities between semantic memory structure and creativity as measured by verbal creativity but not by figural creativity (*38*). Here, we show that individual semantic memory network properties also relate to real-life creativity: individuals with a more compact and less modular organization of their semantic memory exhibit higher creative activities and achievements. This finding is consistent with previous studies reporting a strong relationship between semantic associative ability and creative behavior in real-life (*27*, *28*). It suggests that this relationship may be explained by individual differences in semantic memory structure. We showed that SemNet *modularity* represents both a behavioral marker of real-life creativity and a mediating mechanism underlying the effect of brain functional connectivity on real-life creative activities and achievements. The higher the efficiency and overall connectivity of the brain predictive network, the more flexible the semantic network (characterized by being more compact and less modular), and the more creative the participant is. This result is in line with previous studies (*11*, *36*, *37*) and suggests that more creative individuals have better access to remote concepts within their semantic memory than less creative individuals (*8*, *10*). Importantly, higher modularity in linguistic networks has been linked to rigidity (*67*) and inefficient conceptual processing (*68*). Thus, less modular networks allow more flexible thinking, with a higher connectivity between weakly related elements facilitating their combination.

Previous studies exploring the cognitive processes involved in creativity have revealed brain regions and functional networks associated with different creativity tasks (*5*, *44*, *45*). The use of SemNets allowed us to explore cognitive mechanisms that appear more broadly relevant to associative basis of creative cognition, avoiding the specificities of existing tasks. Using a whole-brain functional connectivity approach, we identified the task-based functional connectivity patterns related to semantic network properties predicting real-life creativity (activities and achievements). These patterns included functional connections distributed across the whole brain, the densest being observed between brain networks previously linked to creativity (*5*, *43*, *53*, *69*–*71*). The major contributions to the prediction of creativity resulted from functional links between control and default mode network, within salience network, and between somatomotor and visual networks. The default mode network has been consistently associated with self-generated thought and spontaneous associations (*16*, *19*, *72*, *73*). In contrast, the control network is associated with controlled processes such as attentional control, working memory, inhibition, memory retrieval, and flexibility, which are necessary to accomplish the objectives of a specific task (*15*, *74*, *75*). The functional coupling between control and default mode networks has been reported in relation to creative cognition in several studies using different approaches such as verbal divergent thinking tasks (*5*, *69*), musical improvisation (*76*), poetry composition (*77*) and visual arts (*78*).

In addition to control and default mode network, the salience network has also been reported to play a critical role in creativity. It has been associated with attentional switching and detection of salient external or internal stimuli and appears to play a role in triggering the engagement of control and default mode networks during creativity tasks (*69*, *79*). Overall, our finding converges with previous correlational (*5*, *69*) and predictive studies of creativity using CPM approach (*43*, *58*) indicating the essential role of the functional connectivity within and between control, default and salience networks for creative thinking abilities.

A considerable number of functional connections between somatomotor and visual networks also contributed to the prediction of creativity via SemNet properties. Both networks have been associated with creativity in previous studies (*53*, *70*, *80*, *81*), but independently. The motor system has been related to creativity (*53*, *82*) as measured by different approaches, including verbal creativity (*71*), music improvisation (*80*, *81*, *83*), and visuospatial creativity (*70*, *71*). The brain regions of visual networks also appear to play an important role mainly in artistic creativity (*71*) and their activation was previously correlated with higher creative achievements (*84*). A recent study using the CPM approach showed the contribution of visual networks in the overlapping brain patterns predicting creativity and intelligence (*58*). Our study adds to this previous work by showing the involvement of the coupling of motor and visual networks in creativity. The role of motor and visual regions in creativity can be plural. In the context of our RJT task used to estimate SemNets, semantic relatedness judgments may evoke visual representations and motor experiences associated with the concepts (*85*). It is then possible that less modular SemNets reflect less segregated motor and visual memory contents in higher creative individuals than in less creative ones, and closer connections between remote concepts in memory.

Overall, our finding further supports and expands existing knowledge on the functional interaction within and between control, default mode and salience networks for creativity (*43*) by showing their link with real-life creativity and characterizing their role in the associative mechanisms captured by SemNet metrics. In addition, the current findings shed light on the contribution of the increased coupling between regions of the visual and motor networks for creativity.

To further characterize the predictive patterns of functional brain connectivity, we identified the nodes with the highest number of connections being localized in the medial prefrontal cortex, insula, the extra-striate inferior region, parietal medial and temporoparietal regions, and temporal pole in the right hemisphere. Most of these regions have been reported to play a role in creative cognition. In a brain lesion study, the medial prefrontal cortex of the default mode network has been shown to be relevant in associative processes underlying creative cognition (*16*). Moreover, this brain region and the insula of the salience network have been highlighted as essential regions for verbal creativity (*5*, *43*, *86*). The right lingual gyrus, part of the extra-striate cortex, is also recruited in verbal creativity tasks (*44*, *45*) in relation to the originality of semantic associations (*28*), and to internally directed attention reflecting increased visual imagery (*87*). Other temporal areas, including the right temporoparietal regions and temporal pole have been associated with verbal and visual creativity (*45*, *88*), including insight problem solving (*89*), and mental imagery (*90*). The involvement of the anterior temporal pole is consistent with its role as a semantic hub (*85*, *91*, *92*) and in abstract thinking and categorization (*93*, *94*).

One surprising result is that the highest degree brain nodes related to real-life creativity were distributed within the right hemisphere. Previous analyses reported a left dominance for creativity regions in functional (*44*, *45*, *50*), connectivity (*43*), and structural (*95*) imaging studies. Most verbal creativity tasks highlight the critical role of brain regions of the left hemisphere, particularly in the prefrontal and temporal cortex, possibly related to linguistic/semantic processing (*44*, *96*, *97*). Here, we also identified left-sided highly connected nodes contributing to the prediction of differences in real-life creativity in the left ventral prefrontal cortex of the control network and in the insula of the salience network, regions that have been shown critical for verbal creativity (*44*, *45*, *98*, *99*). Yet, the right dominance of the predictive patterns in our study was unexpected because our study focused on the semantic basis of creative cognition and used a verbal task. The strong engagement of the right hemisphere might be related to the process of judging remote concepts during the RJT. Previous studies have indeed associated the right hemisphere with a relatively coarser semantic coding (*100*) and the activation of broader semantic fields by words or contexts (*101*). Moreover, the engagement of broad associative processes in the right hemisphere has been related to hemispheric brain asymmetries in dopamine function (*102*). More creative individuals may rate distant words as more related during the RJT than less creative ones, which might rely on a higher functional connectivity with or within the right hemisphere. Hence, these findings show that diverse regions previously reported as central to creative cognition participate together in the predictive connectivity patterns of real-life creativity through a less segregated organization of semantic memory (lower SemNet *modularity*). Whether and how SemNet *modularity* reflects remote thinking that would rely more specifically on the right functional connectivity remain to be addressed in future studies.

Finally, the current SemNets-related results converge with and expand the few recent neuroimaging studies exploring the associative processes of creativity. Higher associative abilities in a free chain association task have been related to higher resting-state functional connectivity within the default mode network (*19*) and to larger gray matter volume in the left posterior inferior temporal gyrus (*49*). In both studies, higher associative abilities mediated the relationship between a priori selected regions of the brain and creativity. One recent study showed that efficiency in SemNets mediated the link between gray matter volume in the left temporal pole and a divergent thinking task (*41*). Our findings advance this knowledge in several critical ways. First, by using SemNets, we were able to estimate the organization of semantic memory, which offers some mechanistic perspective on remote and associative thinking, and showed its role in real-life creativity. Second, we employed a whole brain approach without focusing on a priori regions or networks. Finally, we explored functional connectivity not during rest, but during the RJT, while all participants performed the same trials. This approach minimized individual differences in mental activity during scanning. It importantly gave access to the functional connectivity configuration that occurs during semantic relatedness judgments that reflect semantic associations.

Some limitations to this study need to be acknowledged. First, our sample is relatively small and although the results are robust, the use of additional external validation would add strong support to our findings. Second, we used the SemNet approach that is rooted in the associative theory of creativity (*7*) to estimate individual semantic memory networks based on relatedness judgments of word pairs. The RJT-based SemNet metrics may not capture all the complexity of associative thinking. Thus, future studies are needed to replicate our findings, using alternative methods to estimate individual’s SemNets. How the results generalize across different creative performances and behaviors, in distinct domains, also remains to be explored. Finally, real-life creativity is not exclusively predicted by semantic memory. Many other internal and external factors are important to creativity, such as personality, motivation, emotions and environment (*1*, *2*, *103*–*106*). Despite these other potential dimensions and sources of variability, the brain connectivity patterns allowed us to predict real-life creativity through the individual differences in semantic memory structure, suggesting its strong influence on creative activities and achievements.

In conclusion, the current findings uniquely link brain functional connectivity, semantic memory structure, and real-life creativity by combining advanced network-based methods in novel ways. By exploring semantic memory organization using SemNet methods, we were able to predict creative abilities independently of narrow frameworks or tasks. Our connectome-based modeling approach identified brain connectivity patterns that predicted creative behaviors rooted in semantic memory properties. By converging these two approaches together, our study illustrates how the network organization of the brain and of memory can be related to each other, leading to exciting new frontiers of scientific inquiry.

## Materials and Methods

### Participants

All participants were French native speakers, right-handed, with normal or corrected-to-normal vision and no neurological disorder, cognitive disability or medication affecting the central nervous system. One hundred one healthy participants (48 women) aged between 22 and 40 years (mean 25.6 ± SD 3.7) were recruited via the RISC platform (https://www.risc.cnrs.fr). In total, eight participants were excluded from the fMRI analysis: Six were excluded because of the discovery of MRI brain abnormalities, one fell asleep during the acquisition of the data, and another had a claustrophobia episode at the beginning of the MRI scanning. The latter participant performed the RJT task outside the scanner and was kept in the behavioral analyses only. The final sample was hence composed of 94 participants aged between 22 and 37 years (mean 25.4 ± 4.2) in behavioral analyses and 93 participants in the fMRI analyses (mean age 25.4 ± 3.4; 44 women). A national ethical committee approved the study. After being informed of the study, the participants signed a written consent form. They received monetary compensation for their participation.

### General procedure

Participants underwent a task-based fMRI session during which they performed the Relatedness Judgment Task (RJT). Several training tasks were conducted before acquiring the fMRI data, first outside the scanner, then in the scanner. The training included a motor training task to become familiar with giving responses using the MRI-compatible trackball on a visual scale in the RJT, and a task training to get familiar with the actual task. The task training was similar to the actual task but using different stimuli. In addition, all words used in the RJT were displayed to participants to check that they were familiar with all of them (Details of the task training are described in **SI S1**). After the fMRI session, participants completed a set of creativity tasks on a computer outside the scanner that lasted around three hours.

### Relatedness judgement task (RJT)

#### Task and material description

The RJT has been used to estimate individual-based SemNets and to explore the structure of semantic memory (*36*–*38*). The task requires participants to judge the relatedness of all possible pairs of words from a list of cue words. These judgements are then used to estimate an individuals’ semantic memory network of these words. The selection of the RJT stimuli words used in our study is detailed in Bernard et al. (*37*). In brief, we first created a French SemNet, based on French verbal association norms (*107*) (http://dictaverf.nsu.ru/dictlist), where the nodes represent the words, and the links were weighted by the normative associative strength between words. Next, we computed the shortest path between words and the minimal number of links between each pair was considered as the theoretical semantic distance between the words. Finally, we applied a computational method to select the RJT words that optimized the repartition of the theoretical semantic distance between all possible pairs of these words. The optimal solution included 35 words, resulting in a total of 595 word-pairs that represented the 595 RJT trials.

Each trial began with the displaying word pair on the screen along with a visual scale below ranging from 0 (unrelated) to 100 (strongly related). The stimuli were displayed for 4 seconds in total, divided into a reflection period of 2 seconds to ensure a comparable minimum judgement time and a response period of 2 seconds. During the first two seconds, the participants studied the word pair but couldn’t move the slider yet. Two seconds after stimuli onset, the response period began, the cursor appeared in the middle of the visual scale, and the participants were allowed to move the slider on the visual scale to indicate their rating using a trackball. Participants were instructed to validate their response by clicking the left button of the trackball. The position of the cursor on the scale at the moment of the validation was recorded as the relatedness judgment. When participants did not validate their response, we recorded the slider position at the end of the two-second response period. After the response period, a blank screen was shown during the inter-trial interval jittered from 0.3 to 0.7 seconds (steps = 0.05; **Figure 1a**).

Task trials were distributed into six runs composed of 100 trials each, except for the last run (95 trials). Each run consisted of four blocks of 25 trials each (except the last block of the sixth run with only 20 trials), separated by a 20 second rest period with a cross fixation on the screen. Trials were pseudo-randomly ordered within blocks, such that each block contained a similar proportion of word pairs of each theoretical semantic distance. At the beginning and end of each run, participants had a ten second rest period with a cross fixation on the screen. During the last two seconds of fixation cross periods, the cross changed color, warning the participant that the task was about to start. Participants had a self-paced break inside the scanner between runs.

#### Assessment of individual semantic network structure

##### Building individual semantic networks

The relatedness ratings given by the participant to each pair of words was used to weight the links of the individual SemNet where each word is a node. We represent each of these networks as a 35×35 matrix with one column and one row for each word and cell values correspond to the judgment given by the participant during the RJT task (**Figure 1b**). Based on previous studies and on our pilot study (*36*–*38*), we estimated two types of networks, weighted undirected network (WUN) and unweighted undirected network (UUN; **Figure 1c**). The WUN is a more conservative type of the SemNet, by keeping the weights of all links between the words. The UUN is a less conservative approach, retaining links above a defined threshold, and the links with a weight below the threshold are removed. We defined the threshold as rating value of 50 (the middle of the visual scale) to keep the links between words that were considered moderately or highly associated by the participants. The weights of the remaining links are uniformly transformed to equal 1.

##### Calculation of the individual semantic network metrics

We estimated the properties of the individual SemNet independently for the UUN and the WUN graphs. Based on previous studies relating SemNet to creative abilities (*11*, *34*, *36*–*38*), we computed the following metrics: *ASPL*, *CC*, *Q* and *S* metrics. The Average Shortest Path Length (*ASPL*) is the average shortest number of steps needed to be taken between any pair of nodes. In semantic networks, path length reflects how related two concepts are to each other (*108*, *109*). The Clustering Coefficient (*CC*) measures the network’s connectivity. It refers to the probability that two neighbors of a node will themselves be neighbors. In semantic networks, higher *CC* relates to higher overall relatedness between concepts. Modularity (*Q*) measures how a network is divided (or partitions) into smaller sub-networks; a higher *Q* relates to more sub-communities in the network (*110*, *111*). Such subcommunities can reflect semantic categories in a semantic network. In creativity research, for example, more creative individuals often exhibit a more connected (higher *CC*), less segregated (lower *ASPL* and *Q*) semantic network than less creative individuals ^34^ and these differences were related to flexibility of thought (*35*). The small-worldness (*S*) property of the network is calculated as the ratio between *ASPL* and *CC* and describes how much the nodes that are not directly linked can be reached through connections between their neighbors. In semantic networks, higher *S* has been linked to higher flexibility of thought (*11*). The computations were performed in Matlab, via the Brain Connectivity Toolbox (*112*) (https://www.mathworks.com).

### Assessment of real-life creativity

Outside the scanner, we used the Inventory of Creative Activities and Achievements (*ICAA*) questionnaire (*42*) to assess the real-life creative activities and achievements across eight different creative domains (e.g., literature, music, art and crafts, cooking, sport, visual arts, performing arts, science and engineering). The creative activities (*C-Act*) score reflects the frequency in which participants engaged in various creative activities. Six different questions were posed for each domain, and participants reported the frequency with which they engaged in each activity during the last ten years, using a scale ranging from 0 (never) to 4 (more than ten times). For each participant, the final domain-general score of *C-Act* was the sum of the creative activities across all activities of the eight different domains. The creative achievements (*C-Ach)* score estimated the level of achievement acquired in a creative domain. Ten different levels of achievement were included for each domain going from 0 (never engaged in this domain) to 10 (I have already sold some of my work in this domain). For each participant, the final domain-general score of *C-Ach* was the sum of the scores across the eight different domains.

### Relationships between individual Semantic Network metrics and creativity

We explored whether individual SemNet properties were predictive of real-life creative activities (*C-Act*) and achievements (*C-Ach*; **Figure 1e**). In independent analyses, we performed linear regressions using leave-one-out cross-validations to predict *C-Act* and *C-Ach* scores for each of the SemNet metrics (*ASPL*, *CC*, *Q*, and *S* of WUN and UUN SemNets). The analyses consisted of building a predictive linear model iteratively in N-1 participants using their SemNet metrics (e.g., WUN *Q* SemNet metric) and testing it in the left-out participant. The model was applied on the SemNet metric of the left-out participant to compute a predicted value of the ICAA scores. The significance of the prediction was evaluated via Spearman correlations between the predicted and the observed creativity scores. When the correlations between observed and predicted values were positive with *p* < .05, we assessed its statistical significance using 1,000 iteration permutation testing. We report the Rho coefficient and the *p*-value of the permutation test. Note that Spearman correlations are used for behavioral analyses as creative activities and achievements are typically skewed (*113*). We also ran Spearman correlations between SemNet metrics and ICAA scores to better represent the statistical association between the different SemNet metrics and creativity (**Table 1**).

### MRI Data Acquisition and Preprocessing

Neuroimaging data were acquired on a 3T MRI scanner (Siemens Prisma, Germany) with a 64-channel head coil. Six functional runs were acquired during each six task runs using multi-echo echo-planar imaging (EPI) sequences. No dummy scan was recorded during the acquisition; therefore, we did not discard any volume. Each run included 335 whole-brain volumes acquired with the following parameters: repetition time (TR) = 1,600 ms, echo times (TE) for echo 1 = 15.2 ms, echo 2 = 37.17 ms and echo 3 = 59.14 ms, flip angle = 73°, 54 slices, slice thickness = 2.50 mm, isotropic voxel size 2.5 mm, Ipat acceleration factor = 2, multi-band = 3 and interleaved slice ordering. After the EPI acquisitions, a T1-weighted structural image was acquired with the following parameters: TR = 2,300 ms, TE = 2.76 ms, flip angle = 9°, 192 sagittal slices with a 1 mm thickness, isotropic voxel size 1 mm, Ipat acceleration factor = 2 and interleaved slice order. A resting state fMRI session of 15 minutes followed, not analyzed in the current study.

The preprocessing of the on-task fMRI data was performed for each run separately using the afni_proc.py pipeline from the Analysis of Functional Neuroimages software (AFNI; https://afni.nimh.nih.gov) (*114*). The different preprocessing steps of the data included despiking, slice timing correction and realignment to the first volume (computed on the first echo). We then denoised the preprocessed data using the TE-dependent analysis of multi-echo fMRI data (TEDANA; https://tedana.readthedocs.io/en/stable/), version 0.0.9 (*115*–*117*). The advantage of using multi-echo EPI sequences is that it allows better cleaning of the data by assessing the BOLD and non-BOLD signal through the ICA-based denoising method, improving the reliability of the functional connectivity-based measurement (*118*). The TEDANA pipeline consisted first of an optimal combination of the different echo time series. Then, the dimensionality of the optimally combined data is reduced through the decomposition of the multi-echo BOLD data using principal component analysis (PCA) and independent component analysis (ICA). TEDANA then classifies the resulting components as BOLD or non-BOLD. The exclusion of the non-BOLD components allowed the removal of thermal and physiological noise such as the artefacts generated by the movements, respiration and cardiac activity. The resulting denoised data was co-registered on the T1-weighted structural image using the Statistical Parametric Mapping (SPM) 12 package running in Matlab (Matlab R2017b, The MathWorks, Inc., USA). We then normalized the data to the Montreal Neurological Institute (MNI) template brain, using the transformation matrix computed from the normalization of the T1-weighted structural image, performed with the default settings of the computational anatomy toolbox (CAT 12; http://dbm.neuro.uni-jena.de/cat/) (*119*) implemented in SPM 12. The resulting denoised and normalized images were then entered in a general linear model (GLM) in SPM to covary out the task-related signal from each run. In this analysis, we entered 24 motion parameters (standard motion parameters, first temporal derivatives, standard motion parameters squared and first temporal derivatives squared) and the onsets and durations of each task related events (reflection period, response period, inter trial interval, cross fixation periods and change of the cross-fixation color) as confounds that were regressed from the BOLD signal. We standardized and detrended the residuals of this model for each run and then concatenated the six runs, removing the rest periods between runs (six volumes in total). This final dataset composed of the six task-run residuals concatenated was used as input for the subsequent task-based functional connectivity analyses.

### Building task-based functional connectivity matrices

Calculation of the task-based functional connectivity matrices for each participant was performed using Nilearn v0.3 (*120*) in Python 2.7 (*121*). We used the Schaefer brain atlas to define our ROIs that consisted of 200 ROIs distributed into 17 functional subnetworks than can be summarized in eight main functional networks (*63*). For each ROI, we extracted the BOLD signal during the RJT (averaged across voxels) and computed Pearson correlation coefficients of all pairs of ROIs. As a result, we obtained for each participant a 200×200 matrix with the correlation coefficients between all ROIs. These matrices were Z-Fisher-transform and rescaled in the range of −1 to 1 for the subsequent analyses. This matrix corresponds to the functional connectivity network of each participant in which ROIs are the nodes and correlation coefficients the links.

### A connectome-based predictive modeling approach

We used a CPM approach (*43*, *56*, *57*, *59*) to explore how SemNet properties can be predicted from functional connectivity patterns during the RJT task. We focused the CPM analyses on the SemNet metrics that predicted creativity scores following the method described in Shen et al. (*56*) (**Figure 3**). We used a leave-one-out cross-validation that consisted in building the model iteratively on N-1 participants and test the prediction on the left-out participants.

Since head motions during the fMRI acquisition can affect the CPM results, we verified that there was no correlation between motion patterns during the fMRI acquisition and the SemNet metrics. We estimated the mean FD, that is the sum of the absolute values of the derivatives of the six realignment parameters (*122*), and computed Spearman correlations between the mean FD and all SemNet metrics. The correlations revealed no significant correlation between the motion patterns and WUN *ASPL* (*r* = − .052, *p* = .622), WUN *Q* (*r* = .133, *p* = .203) and UUN *Q* (*r* = .127, *p* = .225).

The first step of the CPM consists of selecting the significant features of brain connectivity to build the “model brain networks”. In the training set (*N*-1), we selected the links of the functional connectivity matrix (correlation coefficients between the ROIs) that significantly correlated with the tested SemNet metric (threshold *p* < .05) either positively (the positive model network) or negatively (the negative model network) across participants (**Figure 3a-b**). Since SemNet metrics had non-Gaussian distributions, we used Spearman correlations. In these model networks of brain connectivity, negative links were removed (*123*). We normalized the values of the links (i.e., the correlation coefficients between ROIs) to have the same range of values for the calculation of the brain networks in the following step.

The second step consists in estimating functional connectivity properties within each participant’s positive and negative model networks. This is one amendment from the classical protocol (*56*) to better take into account the structural properties of functional brain connectivity patterns. Instead of summing the links in the model networks (as in the classical CPM method), we estimated the network properties of the positive and the negative model networks using network metrics (**Figure 3c**). We computed two different whole-brain model network metrics: 1) Network efficiency (*brain-Eff*), measuring rapid and efficient integration across the network (*69*, *124*) and 2) *CC* (*brain-CC*), key property describing a small-world properties network characterizing the human brain (*64*, *65*, *125*–*127*). The *brain-Eff* metric was calculated as the average of the inverse shortest path length. The computation of the *brain-CC* metrics was similar to the *CC* of the SemNet described above in the “Calculation of the individual semantic network metrics” section.

The third and fourth steps consist in building the predictive model using the computed network properties and then applying it to a novel participant (the left out one for each iteration; **Figure 3d**). These steps were conducted separately for each SemNet metric and each model network property. We built a single linear model combining the network metric of the positive and negative model networks of N-1 participants as predictors of a given SemNet metric. The mean FD was included in the model to deal with possible effects of the head motion related to fMRI acquisition on the CPM process. At each iteration, we computed the network metric of the positive and the negative model networks in the left-out participant. We used these values as predictors in the linear model to compute its predicted value of the SemNet metric tested.

The final step evaluated the predictive model by performing a Spearman correlation between the predicted and the observed SemNet metric (*56*). Since we used within-data set cross-validation, for the significant predictions, it was necessary to evaluate the predictive power of the CPM using permutation testing to assess the statistical significance of the results. To this end, we randomly shuffled the values of the SemNet metric 1,000 times, and we ran the new random data through the pipeline of our predictive model in order to generate an empirical null distribution and estimate the distribution of the test statistic given by the correlation between predicted and observed values. The CPM analyses were performed using Matlab Statistical Toolbox (Matlab R2020a, The MathWorks, Inc., USA). The pipeline for the CPM is an adaptation from the protocol by Shen et al. (*56*).

### Functional anatomy of the predicting brain model networks

To explore the patterns of connectivity predicting the SemNet metrics, we characterized the main nodes and links of the significant model networks. We examined the distribution of the connections at the lobar level (between and within brain lobes) and at the intrinsic network level (within and between the eight main functional networks defined by the Schaefer atlas). Finally, we explored the brain distribution of the six highest degree nodes (i.e., ROIs), which are the nodes with the highest number of connections. Due to the nature of the cross-validation approach (running one model for each iteration on N-1 participants), each iteration likely resulted in slightly different links in the model networks. Therefore, we considered the links that were shared between all iterations. The data visualization and plots were performed using BioImage Suite Web 1.0 (http://bisweb.yale.edu/connviewer), BrainNet viewer (*128*) (http://www.nitrc.org/projects/bnv/) in Matlab, and custom scripts in RStudio version 1.3.1056.

### Mediation Analysis

To test whether the patterns of functional connectivity that predict SemNet properties are also relevant for real-life creativity, we ran mediation analyses. For significant CPM predictions, we tested whether the SemNet metrics mediated the relationship between the patterns of brain functional connectivity and creativity. As for the CPM analyses, the mediation analyses focused on the SemNet metrics that correlated with creativity scores. Hence, they explored an indirect effect of the functional brain connectivity on creativity through the SemNet properties.

The mediation analysis (*129*–*131*) consisted in calculating the product of (a) the regression coefficient of the regression analysis on the independent variable (i.e., brain functional connectivity metric, *brain-CC* or *brain-Eff* of the positive or the negative model networks) to predict the mediator (i.e., SemNet metrics) and (b) the regression coefficient of the regression analysis on the mediator to predict the dependent variable (i.e., creativity score), when controlling for the independent variable. We also calculated the regression coefficient of the regression analysis on the independent variable to predict the dependent variable without controlling for the mediator (total effect) and when controlling for it (direct effect; **Figure 6**). All the variables entered in the mediation analyses were normalized, and variables with non-normal distributions were log-transformed. The variables that had a negative correlation with creativity were reversed (multiplied by −1). The selection of the positive or the negative network to be used on the mediation analysis depended on which of them is expected to be positively correlated to the creativity score. We tested the significance of the indirect effect using bootstrapping method, computing unstandardized indirect effects for each 5,000 bootstrapped samples, and the 95% confidence interval was computed by determining the indirect effects at the 2.5^th^ and 97.5^th^ percentiles. The mediation analyses were performed using the PROCESS macro (*132*) in SPSS 22.0 (IBM Corp. in Armonk, NY, USA).

## Acknowledgments

We thank D. Margulies, A. Lopez-Persem, F. De Vico Fallani, M. Chavez for advice and helpful discussion and commentary. We also thank the participants for making this work possible.

## Funding

The research was supported by “Agence Nationale de la Recherche” [grant numbers ANR-19-CE37-0001-01] (EV) and received infrastructure funding from the French programs “Investissements d’avenir” ANR-11-INBS-0006 (EV) and ANR-10-IAIHU-06 (EV). This work was also funded by Becas-Chile of ANID-CONICYT (MOT). The funder had no role in study design, data collection and analysis, decision to publish, or preparation of the manuscript.

## Author Contributions

EV, YNK and MBEN designed the study. MOT, MBER and JB collected the data. MOT analyzed the data with contribution from BB, MBER, TB, JB, EV, and YNK. MOT wrote the first draft of the article. MOT, YNK, MBEN, BB and EV wrote and revised the manuscript. All authors revised and approved the manuscript.

## Competing Interests

The authors declare no competing interests.

## Data and materials availability

The data that support the findings of this study can be available on request from the corresponding author. All the data used in this study were collected on the PRISME and CENIR platforms at the Paris Brain Institute (ICM). Most analyses were conducted using open softwares and toolboxes available online (SPM, AFNI, Nilearn and TEDANA) and using homemade scripts. Custom codes are available from the corresponding authors on request.

## Notes

### Competing Interest Statement

The authors have declared no competing interest.

## References

1. A. Lopez-Persem, T. Bieth, S. Guiet, M. Ovando-Tellez, E. Volle, Through thick and thin: changes in creativity during the first lockdown of the Covid-19 pandemic. (2021), doi:10.31234/osf.io/26qde.

2. T. Lubart, C. Mouchiroud, S. Tordjman, F. Zenasni, Psychologie de la créativité - 2e édition (Armand Colin, Paris, 2e édition., 2015).

3. S. Mastria, S. Agnoli, M. Zanon, T. Lubart, G. E. Corazza, in Exploring Transdisciplinarity in Art and Sciences, Z. Kapoula, E. Volle, J. Renoult, M. Andreatta, Eds. (Springer International Publishing, Cham, 2018; https://doi.org/10.1007/978-3-319-76054-4_1), pp. 3–29.

4. A. Dietrich, The cognitive neuroscience of creativity. Psychon. Bull. Rev. 11, 1011–1026 (2004).

5. R. E. Beaty, M. Benedek, P. J. Silvia, D. L. Schacter, Creative Cognition and Brain Network Dynamics. Trends Cogn. Sci. 20, 87–95 (2016).

6. P. T. Sowden, A. Pringle, L. Gabora, The shifting sands of creative thinking: Connections to dual-process theory. Think. Reason. 21, 40–60 (2015).

7. S. A. Mednick, The associative basis of the creative process. Psychol. Rev. 69, 220–232 (1962).

8. E. Rossmann, A. Fink, Do creative people use shorter associative pathways? Personal. Individ. Differ. 49, 891–895 (2010).

9. R. E. Beaty, P. J. Silvia, E. C. Nusbaum, E. Jauk, M. Benedek, The roles of associative and executive processes in creative cognition. Mem. Cognit. 42, 1186–1197 (2014).

10. M. Benedek, A. C. Neubauer, Revisiting Mednick’s Model on Creativity-Related Differences in Associative Hierarchies. Evidence for a Common Path to Uncommon Thought. J. Creat. Behav. 47, 273–289 (2013).

11. Y. N. Kenett, D. Anaki, M. Faust, Investigating the structure of semantic networks in low and high creative persons. Front. Hum. Neurosci. 8 (2014), doi:10.3389/fnhum.2014.00407.

12. E. Volle, in The Cambridge Handbook of the Neuroscience of Creativity (Editors: R.E. Jung and O. Vartanian, New York:, Cambridge University Press., 2017), Cambridge Handbooks in Psychology.

13. D. Bendetowicz, M. Urbanski, C. Aichelburg, R. Levy, E. Volle, Brain morphometry predicts individual creative potential and the ability to combine remote ideas. Cortex J. Devoted Study Nerv. Syst. Behav. 86, 216–229 (2017).

14. M. Benedek, T. Könen, A. C. Neubauer, Associative abilities underlying creativity. Psychol. Aesthet. Creat. Arts. 6, 273–281 (2012).

15. M. Benedek, E. Jauk, in The Oxford Handbook of Spontaneous Thought: Mind-Wandering, Creativity, and Dreaming (2018).

16. D. Bendetowicz, M. Urbanski, B. Garcin, C. Foulon, R. Levy, M.-L. Bréchemier, C. Rosso, M. Thiebaut de Schotten, E. Volle, Two critical brain networks for generation and combination of remote associations. Brain J. Neurol. 141, 217–233 (2018).

17. T. Paulin, D. Roquet, Y. N. Kenett, G. Savage, M. Irish, The effect of semantic memory degeneration on creative thinking: A voxel-based morphometry analysis. NeuroImage. 220, 117073 (2020).

18. M. P. Ovando-Tellez, T. Bieth, M. Bernard, E. Volle, The contribution of the lesion approach to the neuroscience of creative cognition. Curr. Opin. Behav. Sci. 27, 100–108 (2019).

19. T. R. Marron, E. Berant, V. Axelrod, M. Faust, Spontaneous cognition and its relationship to human creativity: a functional connectivity study involving a chain free association task. NeuroImage, 117064 (2020).

20. C. S. Lee, D. J. Therriault, The cognitive underpinnings of creative thought: A latent variable analysis exploring the roles of intelligence and working memory in three creative thinking processes. Intelligence. 41, 306–320 (2013).

21. R. E. Beaty, D. C. Zeitlen, B. S. Baker, Y. N. Kenett, Forward Flow and Creative Thought: Assessing Associative Cognition and its Role in Divergent Thinking. Think. Ski. Creat., 100859 (2021).

22. M. T. Mednick, S. A. Mednick, C. C. Jung, Continual association as a function of level of creativity and type of verbal stimulus. J. Abnorm. Psychol. 69, 511–515 (1964).

23. A. E. Green, D. J. M. Kraemer, J. A. Fugelsang, J. R. Gray, K. N. Dunbar, Neural correlates of creativity in analogical reasoning. J. Exp. Psychol. Learn. Mem. Cogn. 38, 264–272 (2012).

24. A. E. Green, K. A. Spiegel, E. J. Giangrande, A. B. Weinberger, N. M. Gallagher, P. E. Turkeltaub, Thinking Cap Plus Thinking Zap: tDCS of Frontopolar Cortex Improves Creative Analogical Reasoning and Facilitates Conscious Augmentation of State Creativity in Verb Generation. Cereb. Cortex N. Y. NY. 27, 2628–2639 (2017).

25. T. Merten, I. Fischer, Creativity, personality and word association responses: associative behaviour in forty supposedly creative persons. Personal. Individ. Differ. 27, 933–942 (1999).

26. A. Gruszuka, E. Nęcka, Priming and acceptance of close and remote associations by creative and less creative people. Creat. Res. J. 14, 193–205 (2002).

27. R. Prabhakaran, A. E. Green, J. R. Gray, Thin slices of creativity: Using single-word utterances to assess creative cognition. Behav. Res. Methods. 46, 641–659 (2014).

28. M. Benedek, J. Jurisch, K. Koschutnig, A. Fink, R. E. Beaty, Elements of creative thought: Investigating the cognitive and neural correlates of association and bi-association processes. NeuroImage. 210, 116586 (2020).

29. C. S. Q. Siew, D. U. Wulff, N. M. Beckage, Y. N. Kenett, Cognitive Network Science: A Review of Research on Cognition through the Lens of Network Representations, Processes, and Dynamics. Complexity. 2019, e2108423 (2019).

30. A. Baronchelli, R. Ferrer-i-Cancho, R. Pastor-Satorras, N. Chater, M. H. Christiansen, Networks in cognitive science. Trends Cogn. Sci. 17, 348–360 (2013).

31. J. Borge-Holthoefer, A. Arenas, Semantic Networks: Structure and Dynamics. Entropy. 12, 1264–1302 (2010).

32. J. Borge-Holthoefer, A. Arenas, in Int. j. complex syst. sci. (2011; https://zaguan.unizar.es/record/61329).

33. Y. N. Kenett, in Exploring Transdisciplinarity in Art and Sciences, Z. Kapoula, E. Volle, J. Renoult, M. Andreatta, Eds. (Springer International Publishing, Cham, 2018; https://doi.org/10.1007/978-3-319-76054-4_3), pp. 49–75.

34. Y. N. Kenett, M. Faust, A Semantic Network Cartography of the Creative Mind. Trends Cogn. Sci. 23, 271–274 (2019).

35. A. L. Cosgrove, Y. N. Kenett, R. E. Beaty, M. T. Diaz, Quantifying flexibility in thought: The resiliency of semantic networks differs across the lifespan. Cognition. 211, 104631 (2021).

36. M. Benedek, Y. N. Kenett, K. Umdasch, D. Anaki, M. Faust, A. C. Neubauer, How semantic memory structure and intelligence contribute to creative thought: a network science approach. Think. Reason. 23, 158–183 (2017).

37. M. Bernard, Y. Kenett, M. Ovando-Tellez, M. Benedek, E. Volle, Building Individual Semantic Networks and Exploring their Relationships with Creativity (2019).

38. L. He, Y. N. Kenett, K. Zhuang, C. Liu, R. Zeng, T. Yan, T. Huo, J. Qiu, The relation between semantic memory structure, associative abilities, and verbal and figural creativity. Think. Reason. 27, 268–293 (2021).

39. M. A. Runco, G. J. Jaeger, The Standard Definition of Creativity. Creat. Res. J. 24, 92–96 (2012).

40. S. Acar, M. A. Runco, Divergent thinking: New methods, recent research, and extended theory. Psychol. Aesthet. Creat. Arts. 13, 153–158 (2019).

41. T. Yan, K. Zhuang, L. He, C. Liu, R. Zeng, J. Qiu, Left temporal pole contributes to creative thinking via an individual semantic network. Psychophysiology. e13841 (2021).

42. J. Diedrich, E. Jauk, P. J. Silvia, J. M. Gredlein, A. C. Neubauer, M. Benedek, Assessment of real-life creativity: The Inventory of Creative Activities and Achievements (ICAA). Psychol. Aesthet. Creat. Arts. 12, 304–316 (2018).

43. R. E. Beaty, Y. N. Kenett, A. P. Christensen, M. D. Rosenberg, M. Benedek, Q. Chen, A. Fink, J. Qiu, T. R. Kwapil, M. J. Kane, P. J. Silvia, Robust prediction of individual creative ability from brain functional connectivity. Proc. Natl. Acad. Sci. 115, 1087–1092 (2018).

44. G. Gonen-Yaacovi, L. C. de Souza, R. Levy, M. Urbanski, G. Josse, E. Volle, Rostral and caudal prefrontal contribution to creativity: a meta-analysis of functional imaging data. Front. Hum. Neurosci. 7 (2013), doi:10.3389/fnhum.2013.00465.

45. M. Boccia, L. Piccardi, L. Palermo, R. Nori, M. Palmiero, Where do bright ideas occur in our brain? Meta-analytic evidence from neuroimaging studies of domain-specific creativity. Front. Psychol. 6 (2015), doi:10.3389/fpsyg.2015.01195.

46. M. Benedek, in Exploring Transdisciplinarity in Art and Sciences, Z. Kapoula, E. Volle, J. Renoult, M. Andreatta, Eds. (Springer International Publishing, Cham, 2018; https://doi.org/10.1007/978-3-319-76054-4_2), pp. 31–48.

47. R. E. Beaty, P. Seli, D. L. Schacter, Network Neuroscience of Creative Cognition: Mapping Cognitive Mechanisms and Individual Differences in the Creative Brain. Curr. Opin. Behav. Sci. 27, 22–30 (2019).

48. D. L. Zabelina, J. R. Andrews-Hanna, Dynamic network interactions supporting internally-oriented cognition. Curr. Opin. Neurobiol. 40, 86–93 (2016).

49. C. Liu, Z. Ren, K. Zhuang, L. He, T. Yan, R. Zeng, J. Qiu, Semantic association ability mediates the relationship between brain structure and human creativity. Neuropsychologia. 151, 107722 (2021).

50. L. S. Cogdell-Brooke, P. T. Sowden, I. R. Violante, H. E. Thompson, A meta-analysis of functional magnetic resonance imaging studies of divergent thinking using activation likelihood estimation. Hum. Brain Mapp. 41, 5057–5077 (2020).

51. M. Benedek, T. Schües, R. E. Beaty, E. Jauk, K. Koschutnig, A. Fink, A. C. Neubauer, To create or to recall original ideas: Brain processes associated with the imagination of novel object uses. Cortex. 99, 93–102 (2018).

52. K. Madore, P. Thakral, R. Beaty, D. Addis, D. Schacter, Neural Mechanisms of Episodic Retrieval Support Divergent Creative Thinking. Cereb. Cortex N. Y. N 1991. 29, 1–17 (2017).

53. H. E. Matheson, Y. N. Kenett, The role of the motor system in generating creative thoughts. NeuroImage. 213, 116697 (2020).

54. T. Wei, X. Liang, Y. He, Y. Zang, Z. Han, A. Caramazza, Y. Bi, Predicting Conceptual Processing Capacity from Spontaneous Neuronal Activity of the Left Middle Temporal Gyrus. J. Neurosci. 32, 481–489 (2012).

55. Q. Chen, W. Yang, W. Li, D. Wei, H. Li, Q. Lei, Q. Zhang, J. Qiu, Association of creative achievement with cognitive flexibility by a combined voxel-based morphometry and resting-state functional connectivity study. NeuroImage. 102 Pt 2, 474–483 (2014).

56. X. Shen, E. S. Finn, D. Scheinost, M. D. Rosenberg, M. M. Chun, X. Papademetris, R. T. Constable, Using connectome-based predictive modeling to predict individual behavior from brain connectivity. Nat. Protoc. 12, 506–518 (2017).

57. M. D. Rosenberg, E. S. Finn, D. Scheinost, X. Papademetris, X. Shen, R. T. Constable, M. M. Chun, A neuromarker of sustained attention from whole-brain functional connectivity. Nat. Neurosci. 19, 165–171 (2016).

58. E. Frith, D. B. Elbich, A. P. Christensen, M. D. Rosenberg, Q. Chen, M. J. Kane, P. J. Silvia, P. Seli, R. E. Beaty, Intelligence and creativity share a common cognitive and neural basis. J. Exp. Psychol. Gen. 150, 609–632 (2021).

59. E. V. Goldfarb, M. D. Rosenberg, D. Seo, R. T. Constable, R. Sinha, Hippocampal seed connectome-based modeling predicts the feeling of stress. Nat. Commun. 11, 2650 (2020).

60. Z. Ren, R. J. Daker, L. Shi, J. Sun, R. E. Beaty, X. Wu, Q. Chen, W. Yang, I. M. Lyons, A. E. Green, J. Qiu, Connectome-Based Predictive Modeling of Creativity Anxiety. NeuroImage. 225, 117469 (2021).

61. P. Liu, W. Yang, K. Zhuang, D. Wei, R. Yu, X. Huang, J. Qiu, The functional connectome predicts feeling of stress on regular days and during the COVID-19 pandemic. Neurobiol. Stress. 14, 100285 (2021).

62. M. D. Humphries, K. Gurney, Network ‘Small-World-Ness’: A Quantitative Method for Determining Canonical Network Equivalence. PLOS ONE. 3, e0002051 (2008).

63. A. Schaefer, R. Kong, E. M. Gordon, T. O. Laumann, X.-N. Zuo, A. J. Holmes, S. B. Eickhoff, B. T. T. Yeo, Local-Global Parcellation of the Human Cerebral Cortex from Intrinsic Functional Connectivity MRI. Cereb. Cortex N. Y. N 1991. 28, 3095–3114 (2018).

64. D. J. Watts, S. H. Strogatz, Collective dynamics of ‘small-world’ networks. Nature. 393, 440–442 (1998).

65. O. Sporns, The human connectome: a complex network. Ann. N. Y. Acad. Sci. 1224, 109–125 (2011).

66. F. De Vico Fallani, J. Richiardi, M. Chavez, S. Achard, Graph analysis of functional brain networks: practical issues in translational neuroscience. Philos. Trans. R. Soc. B Biol. Sci. 369, 20130521 (2014).

67. Y. N. Kenett, R. Gold, M. Faust, The hyper-modular associative mind: A computational analysis of associative responses of persons with Asperger syndrome. Lang. Speech. 59, 297–317 (2016).

68. C. S. Q. Siew, Community structure in the phonological network. Front. Psychol. 4 (2013), doi:10.3389/fpsyg.2013.00553.

69. R. E. Beaty, M. Benedek, S. Barry Kaufman, P. J. Silvia, Default and Executive Network Coupling Supports Creative Idea Production. Sci. Rep. 5, 10964 (2015).

70. L. Aziz-Zadeh, S.-L. Liew, F. Dandekar, Exploring the neural correlates of visual creativity. Soc. Cogn. Affect. Neurosci. 8, 475–480 (2013).

71. Q. Chen, R. E. Beaty, Z. Cui, J. Sun, H. He, K. Zhuang, Z. Ren, G. Liu, J. Qiu, Brain hemispheric involvement in visuospatial and verbal divergent thinking. NeuroImage. 202, 116065 (2019).

72. M. E. Raichle, The restless brain: how intrinsic activity organizes brain function. Philos. Trans. R. Soc. Lond. B. Biol. Sci. 370 (2015), doi:10.1098/rstb.2014.0172.

73. T. R. Marron, Y. Lerner, E. Berant, S. Kinreich, I. Shapira-Lichter, T. Hendler, M. Faust, Chain free association, creativity, and the default mode network. Neuropsychologia. 118, 40–58 (2018).

74. R. E. Beaty, A. P. Christensen, M. Benedek, P. J. Silvia, D. L. Schacter, Creative constraints: Brain activity and network dynamics underlying semantic interference during idea production. NeuroImage. 148, 189–196 (2017).

75. E. Mandonnet, M. Vincent, A. Valero-Cabré, V. Facque, M. Barberis, F. Bonnetblanc, F. Rheault, E. Volle, M. Descoteaux, D. S. Margulies, Network-level causal analysis of set-shifting during trail making test part B: A multimodal analysis of a glioma surgery case. Cortex J. Devoted Study Nerv. Syst. Behav. 132, 238–249 (2020).

76. A. L. Pinho, F. Ullén, M. Castelo-Branco, P. Fransson, Ö. de Manzano, Addressing a Paradox: Dual Strategies for Creative Performance in Introspective and Extrospective Networks. Cereb. Cortex. 26, 3052–3063 (2016).

77. S. Liu, M. G. Erkkinen, M. L. Healey, Y. Xu, K. E. Swett, H. M. Chow, A. R. Braun, Brain activity and connectivity during poetry composition: Toward a multidimensional model of the creative process. Hum. Brain Mapp. 36, 3351–3372 (2015).

78. M. Ellamil, C. Dobson, M. Beeman, K. Christoff, Evaluative and generative modes of thought during the creative process. NeuroImage. 59, 1783–1794 (2012).

79. L. Q. Uddin, Salience processing and insular cortical function and dysfunction. Nat. Rev. Neurosci. 16, 55–61 (2015).

80. A. L. Pinho, O. de Manzano, P. Fransson, H. Eriksson, F. Ullen, Connecting to Create: Expertise in Musical Improvisation Is Associated with Increased Functional Connectivity between Premotor and Prefrontal Areas. J. Neurosci. 34, 6156–6163 (2014).

81. R. E. Beaty, The neuroscience of musical improvisation. Neurosci. Biobehav. Rev. 51, 108–117 (2015).

82. Q. Chen, R. E. Beaty, J. Qiu, Mapping the artistic brain: Common and distinct neural activations associated with musical, drawing, and literary creativity. Hum. Brain Mapp. 41, 3403–3419 (2020).

83. C. J. Limb, A. R. Braun, Neural Substrates of Spontaneous Musical Performance: An fMRI Study of Jazz Improvisation. PLoS ONE. 3, e1679 (2008).

84. K. Japardi, S. Bookheimer, K. Knudsen, D. G. Ghahremani, R. M. Bilder, Functional magnetic resonance imaging of divergent and convergent thinking in Big-C creativity. Neuropsychologia. 118, 59–67 (2018).

85. J. R. Binder, R. H. Desai, The neurobiology of semantic memory. Trends Cogn. Sci. 15, 527–536 (2011).

86. R. E. Jung, The structure of creative cognition in the human brain. Front. Hum. Neurosci. 7 (2013), doi:10.3389/fnhum.2013.00330.

87. M. Benedek, in The Cambridge Handbook of the Neuroscience of Creativity, R. E. Jung, O. Vartanian, Eds. (Cambridge University Press, ed. 1, 2018; https://www.cambridge.org/core/product/identifier/9781316556238%23CN-bp-10/type/book_part), pp. 180–194.

88. T. Asari, S. Konishi, K. Jimura, J. Chikazoe, N. Nakamura, Y. Miyashita, Right temporopolar activation associated with unique perception. NeuroImage. 41, 145–152 (2008).

89. L. Aziz-Zadeh, J. T. Kaplan, M. Iacoboni, “Aha!”: The neural correlates of verbal insight solutions. Hum. Brain Mapp. 30, 908–916 (2009).

90. A. Abraham, S. Beudt, D. V. M. Ott, D. Yves von Cramon, Creative cognition and the brain: Dissociations between frontal, parietal–temporal and basal ganglia groups. Brain Res. 1482, 55–70 (2012).

91. M. A. Lambon Ralph, Neurocognitive insights on conceptual knowledge and its breakdown. Philos. Trans. R. Soc. B Biol. Sci. 369, 20120392 (2014).

92. M. A. Lambon Ralph, S. Ehsan, G. A. Baker, T. T. Rogers, Semantic memory is impaired in patients with unilateral anterior temporal lobe resection for temporal lobe epilepsy. Brain. 135, 242–258 (2012).

93. B. Garcin, M. Urbanski, M. Thiebaut de Schotten, R. Levy, E. Volle, Anterior Temporal Lobe Morphometry Predicts Categorization Ability. Front. Hum. Neurosci. 12 (2018), doi:10.3389/fnhum.2018.00036.

94. C. Aichelburg, M. Urbanski, M. Thiebaut de Schotten, F. Humbert, R. Levy, E. Volle, Morphometry of Left Frontal and Temporal Poles Predicts Analogical Reasoning Abilities. Cereb. Cortex. 26, 915–932 (2016).

95. B. Shi, X. Cao, Q. Chen, K. Zhuang, J. Qiu, Different brain structures associated with artistic and scientific creativity: a voxel-based morphometry study. Sci. Rep. 7, 42911 (2017).

96. A. Abraham, K. Pieritz, K. Thybusch, B. Rutter, S. Kröger, J. Schweckendiek, R. Stark, S. Windmann, C. Hermann, Creativity and the brain: Uncovering the neural signature of conceptual expansion. Neuropsychologia. 50, 1906–1917 (2012).

97. A. Abraham, B. Rutter, T. Bantin, C. Hermann, Creative conceptual expansion: A combined fMRI replication and extension study to examine individual differences in creativity. Neuropsychologia. 118, 29–39 (2018).

98. M. Benedek, E. Jauk, A. Fink, K. Koschutnig, G. Reishofer, F. Ebner, A. C. Neubauer, To create or to recall? Neural mechanisms underlying the generation of creative new ideas. NeuroImage. 88, 125–133 (2014).

99. E. G. Chrysikou, H. M. Morrow, A. Flohrschutz, L. Denney, Augmenting ideational fluency in a creativity task across multiple transcranial direct current stimulation montages. Sci. Rep. 11, 8874 (2021).

100. M. Jung-Beeman, Bilateral brain processes for comprehending natural language. Trends Cogn. Sci. 9, 512–518 (2005).

101. C. Chiarello, N. A. Kacinik, C. Shears, S. R. Arambel, L. K. Halderman, C. S. Robinson, Exploring cerebral asymmetries for the verb generation task. Neuropsychology. 20, 88–104 (2006).

102. K. C. Aberg, K. C. Doell, S. Schwartz, The “Creative Right Brain” Revisited: Individual Creativity and Associative Priming in the Right Hemisphere Relate to Hemispheric Asymmetries in Reward Brain Function. Cereb. Cortex. 27, 4946–4959 (2017).

103. T. Lubart, F. Zenasni, B. Barbot, Creative potential and its measurement. Int. J. Talent Dev. Creat. 1, 41–50 (2013).

104. J. A. Plucker, M. Karwowski, J. C. Kaufman, in The Cambridge handbook of intelligence, 2nd ed (Cambridge University Press, New York, NY, US, 2020), pp. 1087–1105.

105. G. J. Feist, A Meta-Analysis of Personality in Scientific and Artistic Creativity. Personal. Soc. Psychol. Rev. 2, 290–309 (1998).

106. M. Karwowski, M. Czerwonka, E. Wiśniewska, B. Forthmann, How Is Intelligence Test Performance Associated with Creative Achievement? A Meta-Analysis. J. Intell. 9, 28 (2021).

107. M. Debrenne, Le dictionnaire des associations verbales du français et ses applications (2011).

108. Y. N. Kenett, E. Levi, D. Anaki, M. Faust, The semantic distance task: Quantifying semantic distance with semantic network path length. J. Exp. Psychol. Learn. Mem. Cogn. 43, 1470–1489 (2017).

109. A. A. Kumar, D. A. Balota, M. Steyvers, Distant connectivity and multiple-step priming in large-scale semantic networks. J. Exp. Psychol. Learn. Mem. Cogn. 46, 2261–2276 (2020).

110. M. E. J. Newman, Modularity and community structure in networks. Proc. Natl. Acad. Sci. 103, 8577–8582 (2006).

111. S. Fortunato, Community detection in graphs. Phys. Rep. 486, 75–174 (2010).

112. M. Rubinov, O. Sporns, Weight-conserving characterization of complex functional brain networks. NeuroImage. 56, 2068–2079 (2011).

113. E. Jauk, M. Benedek, A. C. Neubauer, The Road to Creative Achievement: A Latent Variable Model of Ability and Personality Predictors. Eur. J. Personal. 28, 95–105 (2014).

114. R. W. Cox, AFNI: software for analysis and visualization of functional magnetic resonance neuroimages. Comput. Biomed. Res. Int. J. 29, 162–173 (1996).

115. P. Kundu, N. D. Brenowitz, V. Voon, Y. Worbe, P. E. Vértes, S. J. Inati, Z. S. Saad, P. A. Bandettini, E. T. Bullmore, Integrated strategy for improving functional connectivity mapping using multiecho fMRI. Proc. Natl. Acad. Sci. 110, 16187–16192 (2013).

116. P. Kundu, S. J. Inati, J. W. Evans, W.-M. Luh, P. A. Bandettini, Differentiating BOLD and non-BOLD signals in fMRI time series using multi-echo EPI. NeuroImage. 60, 1759–1770 (2012).

117. The tedana Community, Z. Ahmed, P. A. Bandettini, K. L. Bottenhorn, C. Caballero-Gaudes, L. T. Dowdle, E. DuPre, J. Gonzalez-Castillo, D. Handwerker, S. Heunis, P. Kundu, A. R. Laird, R. Markello, C. J. Markiewicz, T. Maullin-Sapey, S. Moia, T. Salo, I. Staden, J. Teves, E. Uruñuela, M. Vaziri-Pashkam, K. Whitaker, ME-ICA/tedana: 0.0.10 (Zenodo, 2021; https://zenodo.org/record/4725985#.YKjmQus6_RU).

118. C. J. Lynch, J. D. Power, M. A. Scult, M. Dubin, F. M. Gunning, C. Liston, Rapid Precision Functional Mapping of Individuals Using Multi-Echo fMRI. Cell Rep. 33, 108540 (2020).

119. C. Gaser, R. Dahnke, CAT-A Computational Anatomy Toolbox for the Analysis of Structural MRI Data (2016), (available at /paper/CAT-A-Computational-Anatomy-Toolbox-for-the-of-MRI-Gaser-Dahnke/2682c2c5f925da18f465952f1a5c904202ab2693).

120. A. Abraham, F. Pedregosa, M. Eickenberg, P. Gervais, A. Mueller, J. Kossaifi, A. Gramfort, B. Thirion, G. Varoquaux, Machine learning for neuroimaging with scikit-learn. Front. Neuroinformatics. 8 (2014), doi:10.3389/fninf.2014.00014.

121. G. van Rossum, Python reference manual (1995) (available at https://ir.cwi.nl/pub/5008).

122. J. D. Power, A. Mitra, T. O. Laumann, A. Z. Snyder, B. L. Schlaggar, S. E. Petersen, Methods to detect, characterize, and remove motion artifact in resting state fMRI. NeuroImage. 84 (2014), doi:10.1016/j.neuroimage.2013.08.048.

123. A. Fornito, A. Zalesky, M. Breakspear, Graph analysis of the human connectome: promise, progress, and pitfalls. NeuroImage. 80, 426–444 (2013).

124. V. Latora, M. Marchiori, Efficient behavior of small-world networks. Phys. Rev. Lett. 87, 198701 (2001).

125. O. Sporns, C. J. Honey, Small worlds inside big brains. Proc. Natl. Acad. Sci. 103, 19219–19220 (2006).

126. E. Bullmore, O. Sporns, Complex brain networks: graph theoretical analysis of structural and functional systems. Nat. Rev. Neurosci. 10, 186–198 (2009).

127. D. S. Bassett, E. Bullmore, Small-World Brain Networks. The Neuroscientist. 12, 512–523 (2006).

128. M. Xia, J. Wang, Y. He, BrainNet Viewer: A Network Visualization Tool for Human Brain Connectomics. PLoS ONE. 8, e68910 (2013).

129. K. J. Preacher, A. F. Hayes, SPSS and SAS procedures for estimating indirect effects in simple mediation models. Behav. Res. Methods Instrum. Comput. 36, 717–731 (2004).

130. A. F. Hayes, Beyond Baron and Kenny: Statistical Mediation Analysis in the New Millennium. Commun. Monogr. 76, 408–420 (2009).

131. D. D. Rucker, K. J. Preacher, Z. L. Tormala, R. E. Petty, Mediation Analysis in Social Psychology: Current Practices and New Recommendations: Mediation Analysis in Social Psychology. Soc. Personal. Psychol. Compass. 5, 359–371 (2011).

132. A. F. Hayes, A. K. Montoya, A Tutorial on Testing, Visualizing, and Probing an Interaction Involving a Multicategorical Variable in Linear Regression Analysis. Commun. Methods Meas. 11, 1–30 (2017).

