## Supplementary material for "Brain connectivity-based prediction of real-life creativity is mediated by semantic memory structure": SI S1, SI Figure 1

**Supplementary text**

**Material and Methods.**

**S1: RJT motor and task training.**

Before the RJT task, participants performed a motor and task training outside the MRI scanner. The motor training was performed for the participants to become familiar with the trackball used as an fMRI response device. The training consisted of 25 trials that had the same structure and timing as in the actual task (**Figure 1a**), but instead of word pairs, a number was presented on the screen. Participants were instructed to locate the proposed number on the visual scale by moving the slider using the trackball. Each training trial began with the display of a random number in the center of the screen along with a visual scale below going from 0 to 100. Numbers could have any of the values of the visual scale, and only the extreme values (0 and 100) were displayed on the visual scale. The stimuli were displayed for 4 seconds in total including a reflection period of 2 seconds, and a response period of 2 seconds. During reflection period, the participants could visualize the position of the proposed number on the visual scale, but they couldn't move the slider yet. During the response period, the cursor appeared in the middle of the visual scale and the participants were allowed to move the slider on the visual scale to locate it on the position of the proposed number. They were asked to validate their response by clicking the left button of the trackball. After the validation, participants received a feedback on the screen indicating the position of the slider on the visual scale at the moment they validated their response. When participants did not validate their response, they received a feedback indicating that they were unable to complete the trial during the response period. At the end of each trial, an inter trial interval jittered from 0.3 to 0.7 seconds (interval = 0.05) occurred before the next trial.

The task training was performed to familiarize the participants with the actual task. This training followed the same principles and timing than the actual task and only varied from it regarding the words used. Participants performed 15 trials. Each trial included the reflection period, the response period and an inter trial interval. Before starting the fMRI acquisition, the participants performed the motor and the task training inside the scanner. In addition, we ensured that all participants were familiar with the 35 words used in the RJT.

Figure S1

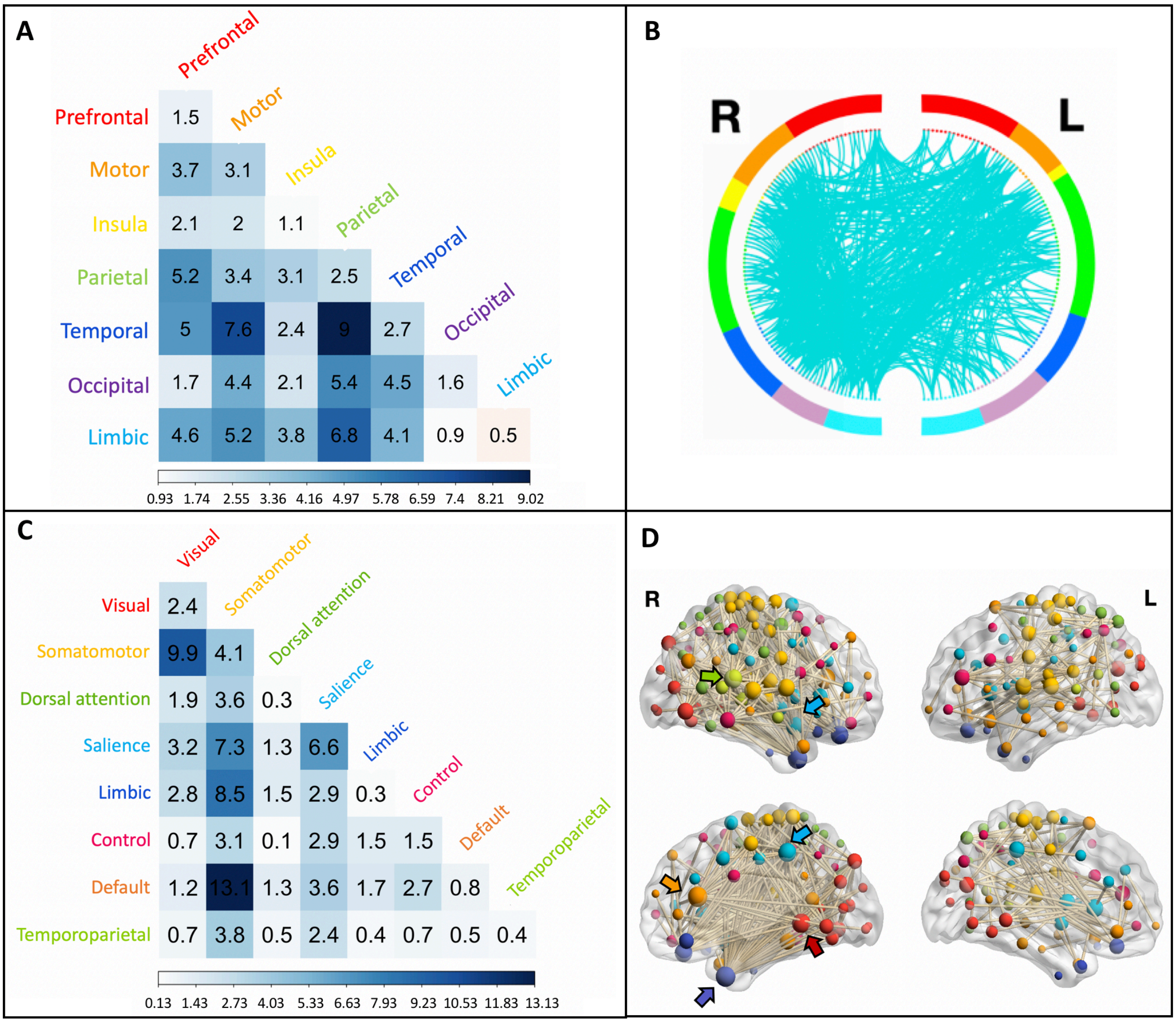


**Supplementary Figure S1. Functional anatomy of the CPM-predicted SemNet metric WUN *Q*.** (**A**) First, we examined the distribution of the links of the model network at the brain location level, specifically into the brain lobes. The correlation matrix represents the percentage of links within the model network connecting seven different brain lobes (total of links = 754). (**B**) A circular graph represents the distribution of links within and between brain regions in the left and right hemispheres. Brain regions are color-coded as in (**A**) and the cyan lines represent the links connecting the ROIs. For visualization purposes, we used a nodal degree threshold of $k = 20$. (**C**) Second, we examined the distribution of the links across intrinsic functional networks based on Schaefer's atlas (*63)*. The matrix represents the percentage of links within the model network occurring within and between eight intrinsic brain networks. (**D**) The nodes and links of the model network are superimposed on a volume rendering of the brain. The color of the nodes represents the functional network they belong to, using a similar color code as in (**B**). The size of the nodes is proportional to their degree, and the highest degree nodes are marked by arrows. Nodes with degree $k = 0$ are not displayed. The highest degree nodes were found in the right hemisphere in the temporal pole (purple arrow, $k = 65$), medial prefrontal cortex (orange arrow, $k = 64$), extra-striate cortex (red arrow, $k = 48$), temporal parietal (green arrow, $k = 44$), parietal medial (light blue arrow, $k = 40$) and insula (light blue arrow, $k = 37$).
